# A stress-dependent postembryonic role for the core CPA factor CFIM-1 in germline integrity

**DOI:** 10.1101/2025.07.01.662614

**Authors:** Anson Sathaseevan, Rhea Ahluwalia, Bin Yu, Claudia Makhenko-Tang, Haruka Nishimura, Alexander Leong, Ka Nam Naomi Kwan, W. Brent Derry

**Affiliations:** Developmental, Stem Cell & Cancer Biology Program, Peter Gilgan Centre for Research and Learning, The Hospital for Sick Children, Toronto, M5G 0A4, ON, Canada; Department of Molecular Genetics, University of Toronto, Toronto, M5S 1A8, ON, Canada; School of Medicine, Queen’s University, Kingston, K7L 3N6, ON, Canada

**Keywords:** CFIM-1, Caenorhabditis elegans, alternative polyadenylation, cleavage and polyadenylation

## Abstract

Post-transcriptional processing of pre-mRNAs generates a diversity of 3’ UTR transcript isoforms that can vary in their function and stability. The differential enrichment of transcript isoforms has been implicated in diseases ranging from cancer to neurodevelopmental disorders. However, the post-embryonic developmental roles of the core ensemble of cleavage and polyadenylation (CPA) factors that mediate these post-transcriptional changes remain poorly characterized. Here, we report a stress-dependent role for the core CPA factor CFIM-1 in germline integrity in *Caenorhabditis elegans.* Total loss-of-function of *cfim-1* elicits a temperature-sensitive sterile phenotype in hermaphrodites. These changes in brood size are accompanied by broad sperm, oocyte and germline morphology defects. Surveying the transcriptome of *cfim-1(lf)* worms, we uncover changes in transcript isoform abundance for dozens of genes with known functions related to the development and maintenance of these structures, consistent with a model in which post-transcriptional regulation of target genes via *cfim- 1* is crucial to the development and maintenance of germline integrity. Our findings collectively define a novel post-embryonic role for a core CPA factor in tissue-specific development.

## INTRODUCTION

The development and function of the hermaphrodite *Caenorhabditis elegans* germline are tightly coordinated processes, relying on a complex interplay among signalling pathways and the regulation of their gene products. The germline is specified early in embryogenesis (Strome 2005; Hubbard & Greenstein, 2005). Following hatching, the gonadal primordium of first stage (L1) larvae contains several primordial germ cells flanked by somatic germ cells. Several rounds of cell division give rise to the mitotic niche that maintains germ cell proliferation. Progression through the larval stages involves several landmark morphological and functional changes, including clear spatial demarcation of the somatic gonad, rapid anatomical extension of the gonad arms, meiotic differentiation, and the initiation of germline sex determination. In the germline, transition to the fourth larval stage (L4) is marked by spermatogenesis, during which a finite pool of spermatids are generated that ultimately determine brood size. The switch to oogenesis occurs in early adulthood and maintains the production of female gametes throughout the life of the organism (Ellis and Schedl, 2007). Ovulation pushes spermatids into the spermatheca, where they mature into competent spermatozoa (Ward and Carrel, 1979; L’Hernault, 2006). During ovulation, oocytes at the proximal germline arm are queued and sequentially passed through the spermatheca where they are fertilized to initiate embryogenesis. In adulthood, mitotically proliferating germ cell nuclei migrate away from the mitotic niche and undergo meiosis before maturing into oocytes that are sequentially queued at the proximal germline. Fertilization proceeds in a conveyor belt-like fashion, beginning with the most proximal oocyte, whereby oocytes at the proximal germline arm are queued and sequentially pushed through the spermatheca. Among the myriad of signalling networks that tightly coordinate development and function of the germline, 3’ UTR based regulation of germline-expressed transcripts via trans-acting factors has been identified as crucial. (Merrit et al., 2008; Marin and Evans, 2003; Kimble & Crittenden, 2005; Hunter & Kenyon, 1996; Wickens et al., 2002; Ogura et al., 2003; Mootz et al., 2004; Mangone et al., 2010). However, little attention has been paid to how 3’ UTR lengths are regulated and how this impacts germline development.

The core ensemble of cleavage and polyadenylation factors facilitate 3’ UTR cleavage site selection and maturation of pre-mRNAs (Steber et al., 2019; Mitschka and Mayr, 2022; **Fig. S2A**). The *cfim-1* gene encodes a component of the cleavage factor Im (CFIm) complex that canonically activates pre-mRNA 3’-end CPA processing and favours distal 3’ UTR site selection (Yang et al., 2010; Yang et al., 2011; Hardy and Norbury, 2016; Brumbaugh et al., 2018). Studies of the orthologs of *cfim-1* across mammalian models have implicated this core CPA factor in diseases ranging from cancer to neurodevelopmental disorders (Alcott et al., 2020; Masamha et al., 2014; Sun et al., 2017; Tan et al., 2018; Chu et al., 2019; Yang et al., 2020; Xing et al., 2021). However, due to its essentiality in this context, the post-embryonic roles, if any, of total loss-of-function of this core CPA factor, and indeed most core CPA factors, remain nebulous (Koscielny et al., 2014).

We previously generated worms with homozygous, complete loss-of-function of *cfim-1* via CRISPR/Cas9 and showed that *cfim-1* is a key regulator of oncogenic Ras/MAPK signaling in the context of somatic vulval development (Subramanian et al., 2021). These findings led us to investigate its role in the function of other tissues, including the germline. Although we found that total loss of *cfim-1* causes defects in fertility, its role in the germline has not been investigated.

Here, we show that *cfim-1(lf)* mutants exhibit a temperature-dependent sterile phenotype that is associated with reproducible morphological changes to the germline as well as defects in spermatogenesis, sperm and oocyte function. Intriguingly, we find that among core CPA factors, these stress-dependent phenotypes are unique to *cfim-1*. Data mining of an RNA sequencing dataset of *cfim-1(lf)* for changes in transcript isoform abundance reveals an enrichment of genes involved in reproductive functions, and also suggests a novel inhibitory model of CPA site selection at 3’ UTRs of pre-mRNAs via CFIM-1 that is distinct from other metazoans.

## RESULTS

### *cfim-1(lf)* elicits temperature-sensitive sterility

We previously observed fecundity defects in the context of total loss-of-function *(lf)* of *cfim-1* (Subramanian et al., 2021). This finding, combined with the robust expression of *cfim-1* in the germline, led us to investigate its role in fertility (**Fig. S1A**). Quantification of the fecundity defect revealed reduced brood sizes relative to wildtype worms at 20 °C (**Fig. 1A**). This effect is exacerbated at a higher temperature, as *cfim-1(lf)* mutants were completely sterile at 25 °C. We validated this sterile phenotype using RNA interference (RNAi) and a separate transgenic line that completely knocks out the open reading frame (ORF) corresponding to *cfim-1* (**Fig. S1C**, **S2B**).

**Figure 1:**
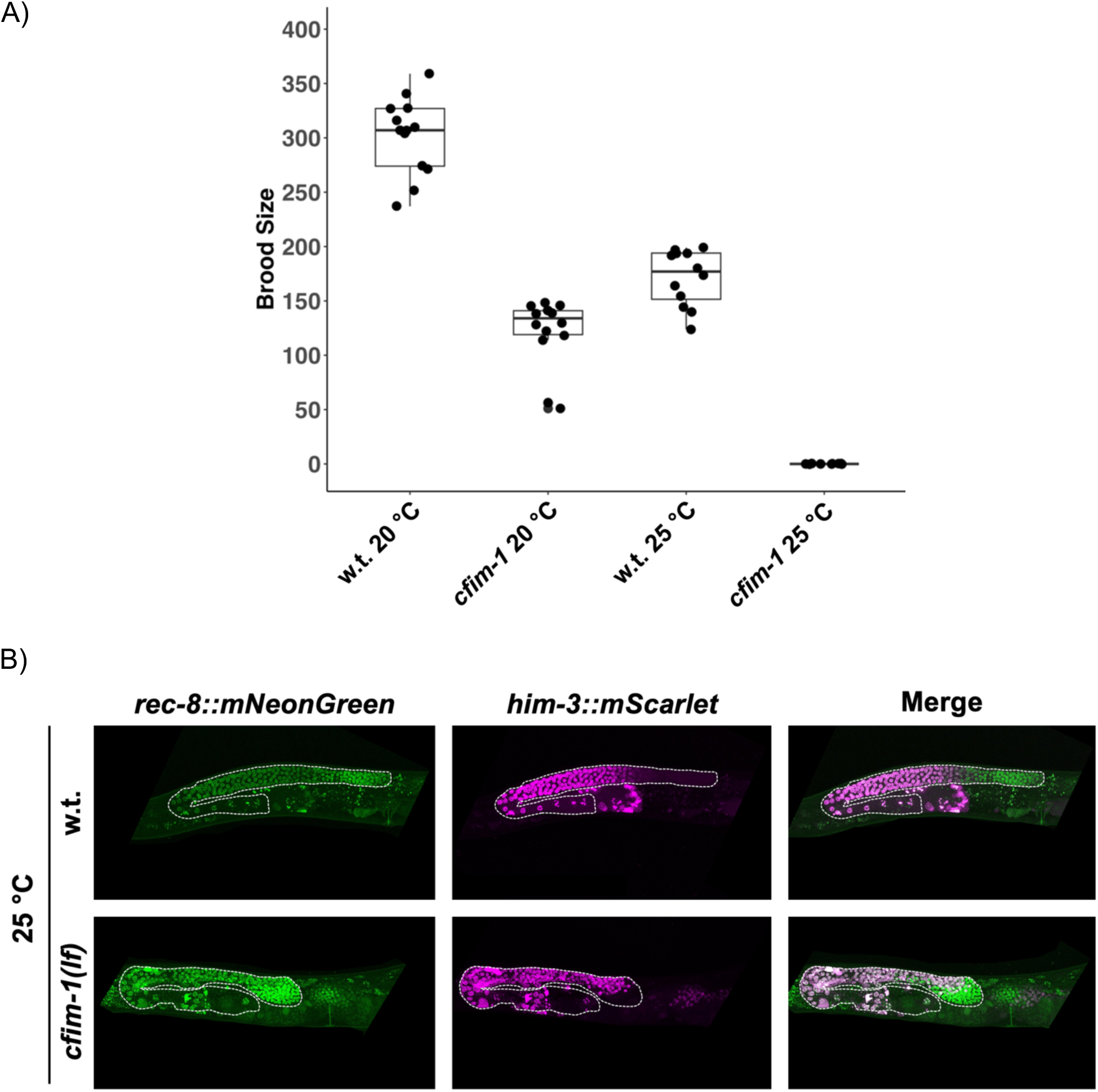
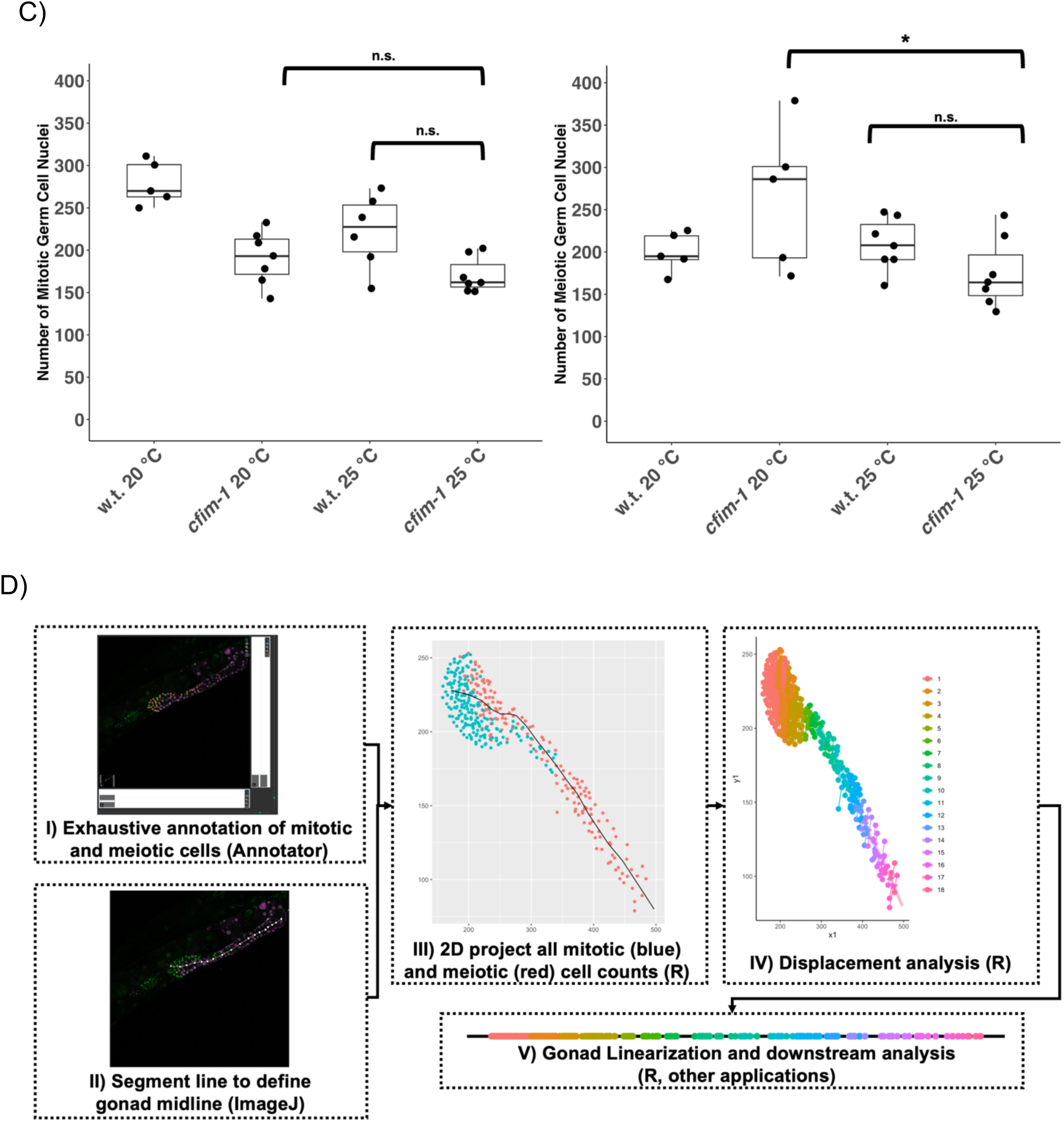
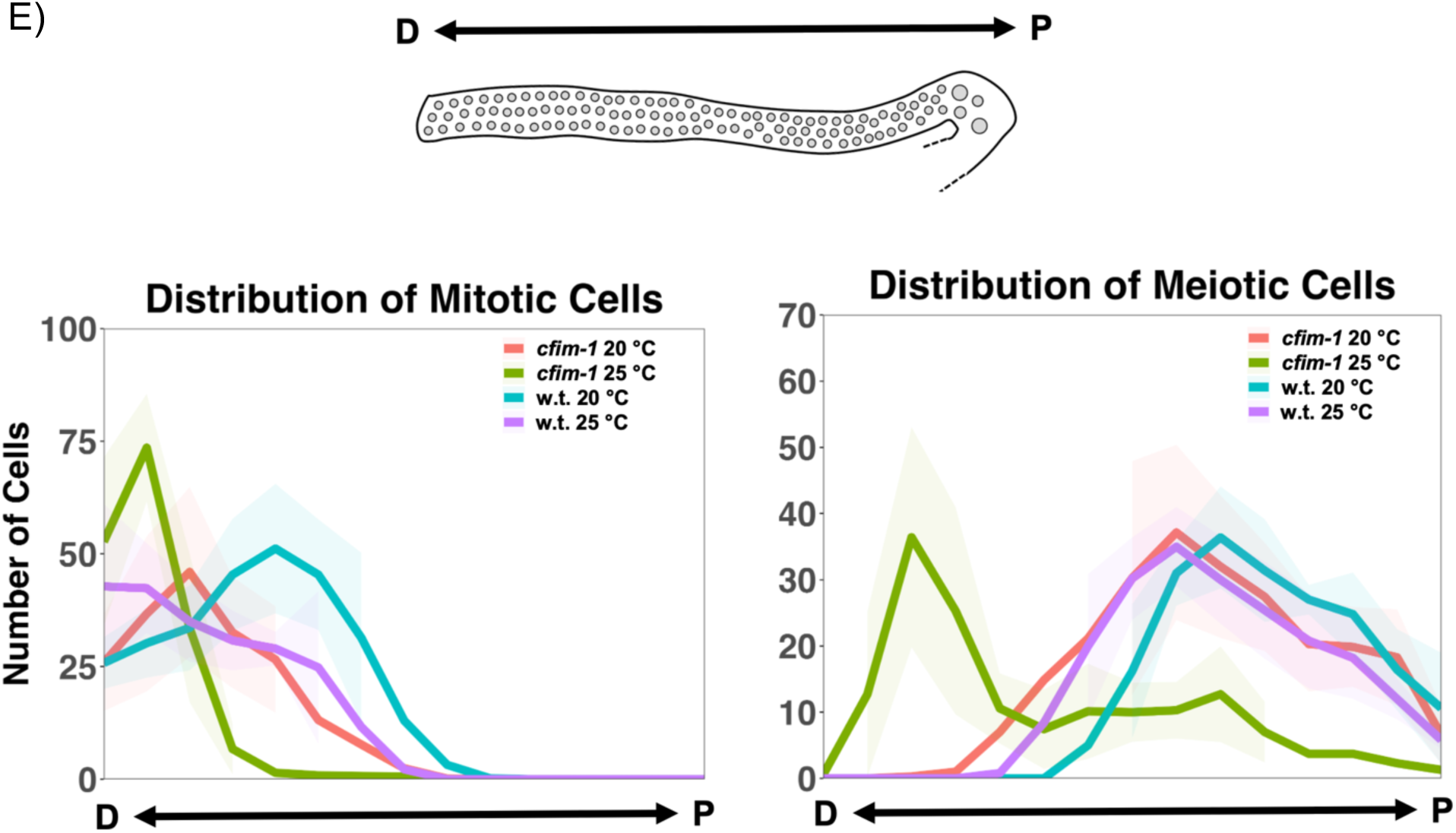
*cfim-1* elicits temperature-dependent sterility with associated germline morphological changes. **A)** Brood counts for *cfim-1(lf)* and N2 wildtype (“w.t.”) worms at the indicated temperatures. Each dot represents the brood size for one worm. Box and whisker overlays show the median with the first and third quartiles. n=13 for wildtype 20 °C, n=14 for *cfim-1* 20 °C, n=12 for wildtype 25 °C and n=10 for *cfim-1 25 °C.* **B)** Representative fluorescence germline images of *cfim-1(lf)* and wildtype worms. Maximum intensity projections are shown with germline (between distal end and terminal oocyte) outlined. **C)** Manual quantifications for the number of mitotic (*rec- 8::mNeonGreen*) and meiotic (*him-3::mScarlet*) germ cell nuclei. Each dot represents one annotated germline. n.s p > 0.05 (Mann-Whitney test); * p < 0.05 (Mann-Whitney test). **D)** Germline annotation and linearization pipeline. **E)** Distribution of mitotic or meiotic cells as a function of position along the distal-proximal (D-P) gonad arm. Lines indicate median values with shaded areas showing standard deviation.

To understand the anatomical basis of this temperature-sensitive sterile phenotype, we examined the organization of germ cell nuclei in *cfim-1* mutants, reasoning that the sterile phenotype might result from proliferative defects in this tissue. To this end, we generated a stable transgenic line that fluorescently tags the endogenous loci of *rec-8* and *him-3*, which are well-established markers of germline mitotic and meiotic nuclei, respectively (Paiserbek et al., 2001; Zetka et al., 1999; Hubbard and Schedl, 2019). While we observed accumulation of germ cell nuclei in the distal gonad of *cfim-1(lf)* mutants, we did not observe dramatic differences in the quantity of germ cell nuclei when reared at 25 °C (**Fig. 1B, C**). Since spatial organization of the germline is important for cell cycle progression, mRNA localization and translation, we wondered if the defects in fecundity were associated with changes in the organization of germ cell nuclei (West et al., 2018; Diag et al., 2018). We used a modified version of an image analysis linearization algorithm that allows for cross-comparison of the distribution of germ cell nuclei in different conditions to test this hypothesis (Toraason et al., 2021; Venkatachalam et al., 2016) (**Fig. 1D**). This approach showed that *cfim-1(lf)* mutants accumulate both mitotic and meiotic germ cell nuclei distally at 25 °C (**Fig. 1E**). However, applying this germline imaging method to several other core CPA factors did not show simultaneous distal accumulation of mitotic and meiotic germ cell nuclei (**Fig. S2E**). We also examined fecundity phenotypes using RNAi to abrogate the function of several core CPA factors and found that none of the other surveyed factors showed a consistent temperature-sensitive decrease in fecundity (**Fig. S2B, D**). Taken together, these results suggest a unique role for CFIM-1 as a core CPA factor in the development and maintenance of gross germline organization and function.

### *cfim-1(lf)* mutants are defective in spermatogenesis

To determine the onset of this sterile phenotype of *cfim-1(lf)* worms reared from hatching at 25 °C, we performed temperature shift experiments from 20 °C to 25 °C at different stages of larval development and quantified broods. We did not notice appreciable changes in broods during temperature shift experiments at the various stages of larval development (**Fig. 2A**). Spermatogenesis is initiated at the L4 stage, followed by a switch to oogenesis when the worms enter adulthood approximately 12 hours later (Ward and Carrel, 1979; Kimble and White, 1981; Kuwabara and Parry, 2001; Pepper et al., 2003). Because the pool of sperm is finite, the brood size of wildtype hermaphrodites is limited to about 300 self-progeny. We hypothesized that the reduced broods of *cfim-1(lf)* mutants might be attributed to compromised spermatogenesis, noting that there are no obvious proliferation defects in mitotic and meiotic germ cell nuclei bound for oocyte fates in mature *cfim-1(lf)* germlines (**Fig. 1C**). Thus, we repeated the temperature shift experiments and quantified broods between the onset of L4 and adulthood (**Fig. 2B**). Shifting worms to 25 °C at progressive timepoints between the onset of L4 and just before egg laying resulted in larger brood sizes in *cfim-1(lf)* mutants, while having no drastic effect on wildtype brood sizes (**Fig. 2B**). Correspondingly, the reciprocal temperature shift experiments from 25 °C to 20 °C resulted in decreased broods in the *cfim-1(lf)* mutants but did not greatly affect wildtype brood sizes.

**Figure 2:**
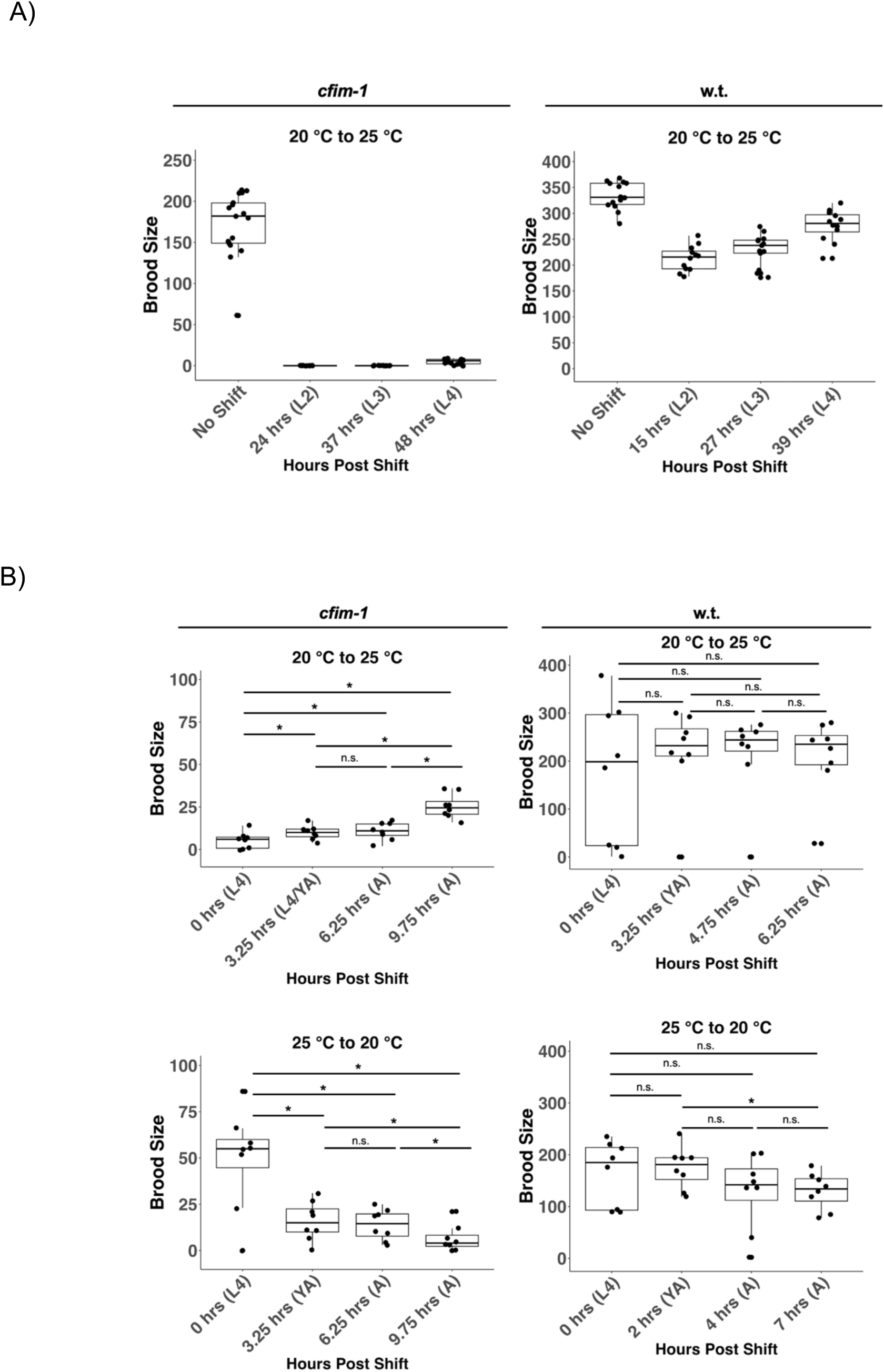
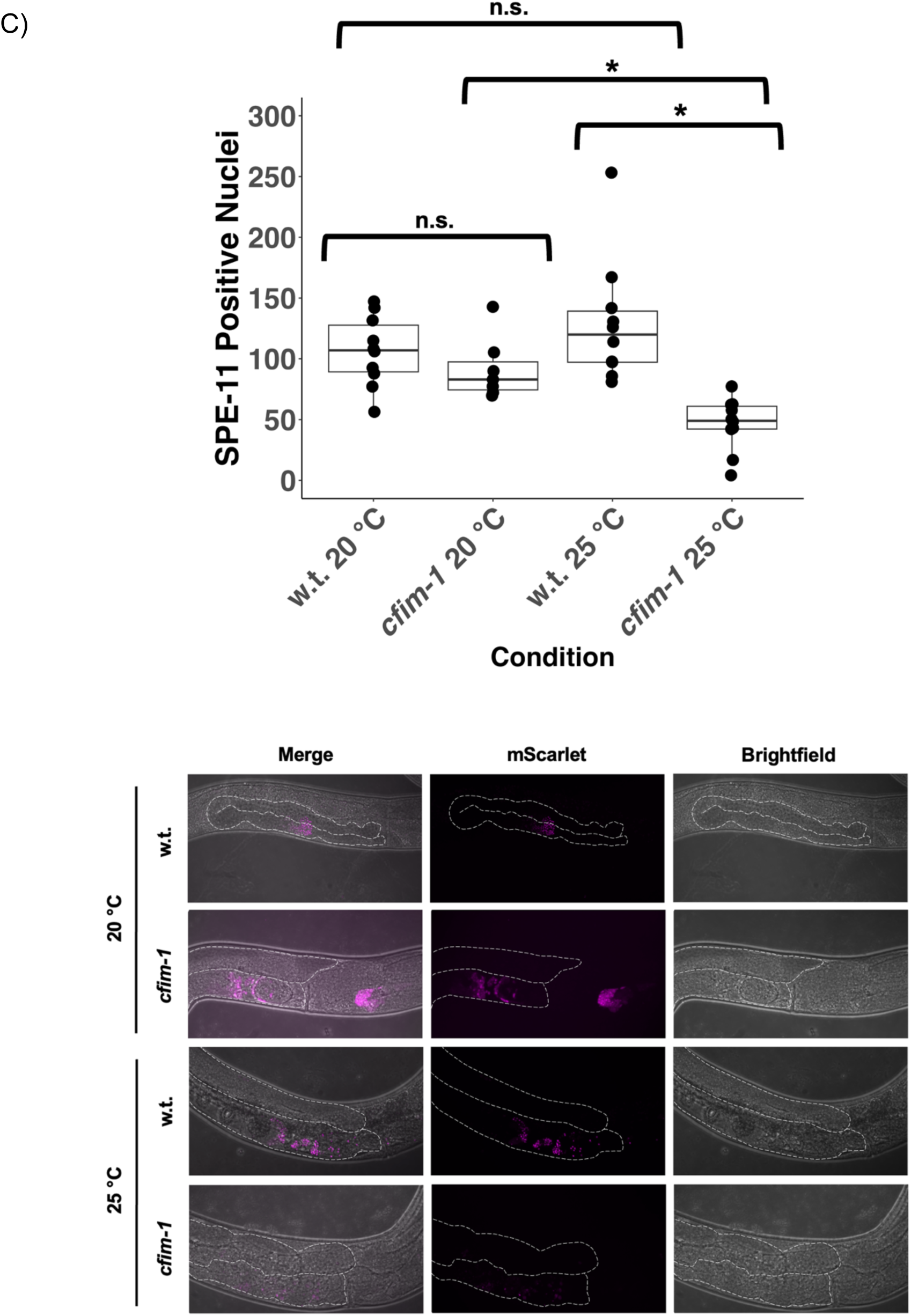
*cfim-1(lf)* worms are defective in spermatogenesis. **A)** Brood sizes of wildtype and *cfim-1(lf)* mutants during temperature shift assays from 20 °C to 25 °C during larval development. **B)** Temperature shift experiments between L4 and adult stages showing brood size quantifications for *cfim-1(lf)* and wildtype (w.t.) worms. Each dot represents the brood size of one worm. n.s p > 0.05 (Mann-Whitney test); * p < 0.05 (Mann-Whitney test). **C)** Quantification of *spe-11::mScarlet* positive nuclei with accompanying representative DIC and fluorescence germline images. Each dot represents the number of spe-11 positive nuclei from the intact gonad of a unique worm. n.s p > 0.05 (Mann-Whitney test); * p < 0.05 (Mann-Whitney test).

To test the hypothesis that sterility of *cfim-1* mutants is due to defects in spermatogenesis, we generated a transgenic line that fluorescently tags the sperm marker SPE-11. In *cfim-1(lf)* mutants reared at the sterility inducing temperature, germ cell nuclei positive for SPE-11 were significantly reduced (**Fig. 2C**). We also noted differences in the distribution of expression of SPE-11 in the germline. In contrast to the punctate, well-demarcated SPE-11::mScarlet positive nuclei in wildtype worms, *cfim-1* mutants displayed diffuse expression in SPE-11::scarlet positive nuclei that have irregularly-defined borders (**Fig. 2C**).

We also examined the distribution of germ cell nuclei positive for SPE-11. Sperm were typically localized between the most recently fertilized embryo and the spermatheca in adult wildtype worms. However, in *cfim-1(lf)* hermaphrodites, sperm were distributed beyond this region, suggesting motility defects (**Fig. 2C, 3A**). We also attempted to evaluate sperm motility by crossing *cfim-1(lf); spe-11::mScarlet* males to both wildtype and *cfim-1(lf)* hermaphrodites at 25 °C. Although these males copulate, they fail to inseminate hermaphrodites. Taken together, these findings collectively suggest that the sperm of *cfim-1(lf)* worms reared at 25 °C are motility-defective.

Recognizing that *cfim-1(lf)* oocytes are morphologically abnormal with variable sizes and relative positions in the proximal gonad at 25 °C (**Fig. S1B**), we also investigated the role of the proximal hermaphrodite tract in facilitating fertilization by mating *spe-11::mScarlet* males to *cfim-1(lf)* and wildtype hermaphrodites at 25 °C. In *cfim-1(lf)* hermaphrodites, sperm aggregate in a spermatheca-like structure, poised to fertilize the queued oocyte. However, the queued oocyte remained unfertilized following ovulation, suggesting oocyte defects (**Video 1, Fig. 3B**). Using a modified version of a sperm motility assay, we also quantified sperm trajectories of germ cell nuclei expressing SPE-11 in the reproductive tract (Hu et al., 2019). On aggregate, the migration speed of sperm were slower in the *cfim-1(lf)* reproductive tracts (**Fig. 3C**).

**Figure 3:**
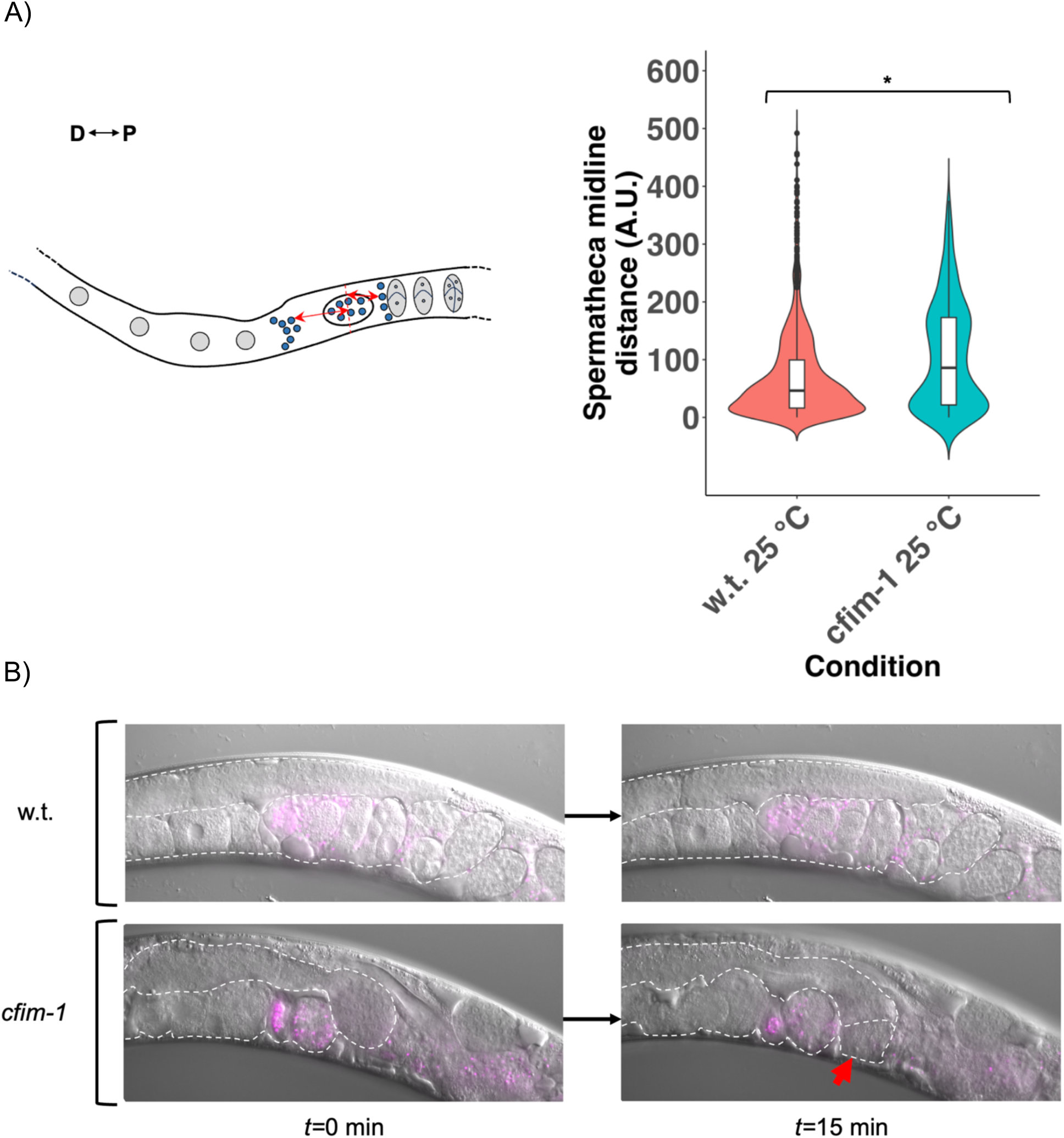
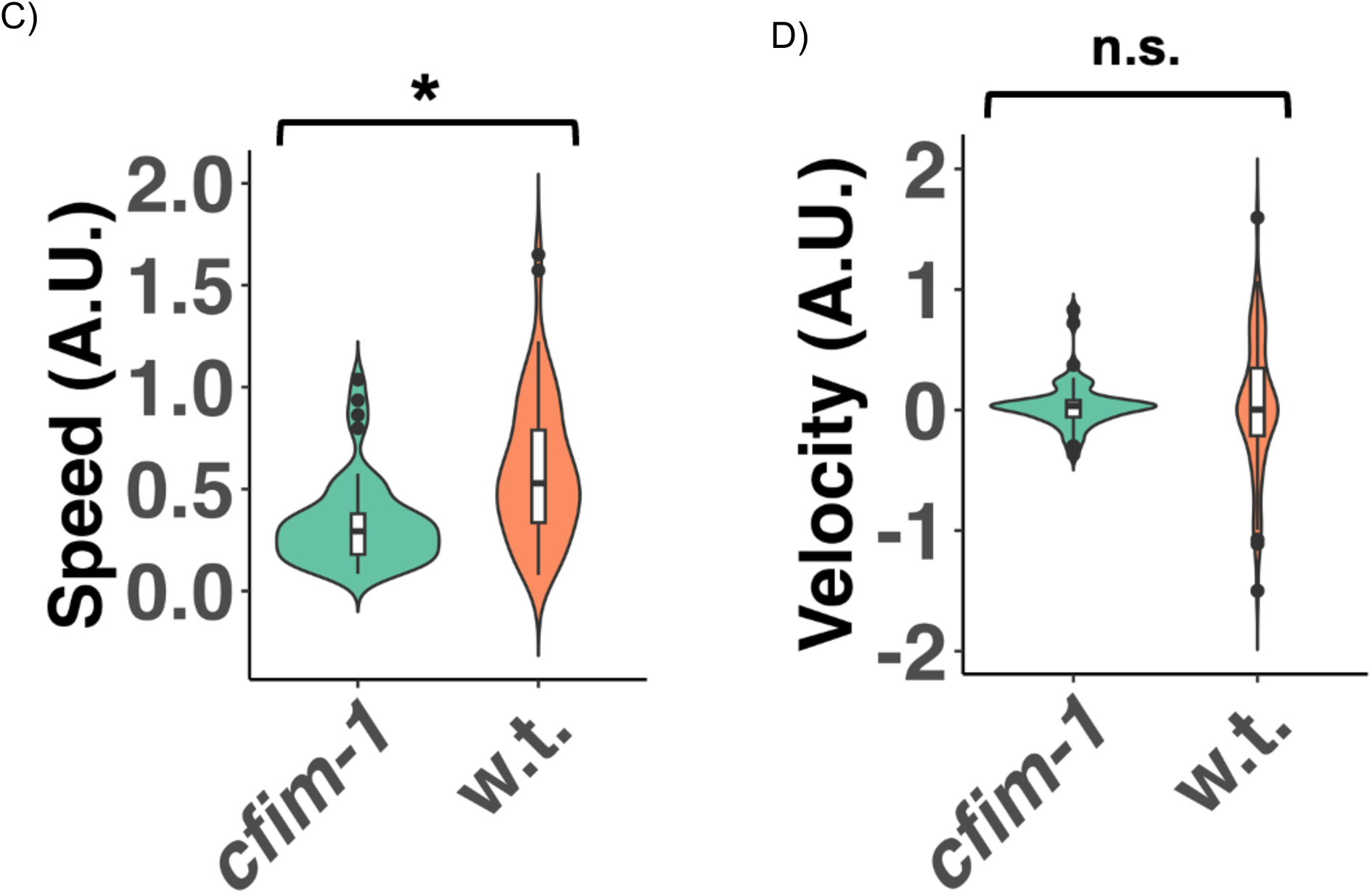
*cfim-1(lf)* mutants exhibit sperm and oocyte functional defects. **A)** Aggregate distance (red arrows in cartoon at left) of sperm from spermatheca midline (red dashed line in cartoon at left) for *cfim-1(lf)* and w.t. worms reared at 25 °C. Each violin plot is the aggregate of sperm distributions for all assayed worms of the indicated genotypes. * p < 0.05 (Kolmogorov-Smirnov test). **B)** Representative merged Nomarski/fluorescence timelapse images of wildtype and *cfim-1* hermaphrodites reared at 25 °C that were mated to *spe-11::mScarlet* males. For the *cfim-1(lf)* images, the red arrow points to an unfertilized oocyte. **C)** Aggregate speed of sperm tracked in the proximal gonad between the vulva and the spermatheca. Each violin represents the aggregate speed of sperm of all mated hermaphrodites of the indicated genotype reared at 25 °C. n.s, p > 0.05 (Mann-Whitney test); * p < 0.05 (Mann-Whitney test). **D)** Velocity of sperm tracked in the proximal gonad between the vulva and the spermatheca. n.s p > 0.05 (Mann-Whitney test); * p < 0.05 (Mann-Whitney test)

While we did not note significant changes in vector velocity of the sperm relative to the spermatheca in wildtype versus *cfim-1(lf)* worms, we did observe a smaller spread of velocity values in *cfim-1(lf)* gonads (**Fig. 3D**). This may reflect constraints in terms of the ability of these sperm to navigate the proximal gonad tract.

### Fertility genes are subject to APA regulation by CFIM-1

Given the known roles of CFIM-1 in 3’ end processing in the regulation of hundreds of genes and our observations that spermatogenesis is compromised in *cfim-1(lf)* mutants, we surveyed the transcriptome of *cfim-1(lf)* mutants (Subramanian et al., 2021). Along with others, we previously demonstrated that loss of function of *cfim-1* and its mammalian homolog, *NUDT21*, results in global biasing towards utilization of proximal cleavage sites of the 3’ UTRs of pre-mRNAs bound for transcriptional processing, a process referred to as “3’ UTR shortening” (Masamha et al., 2014; Brumbaugh et al., 2018; Subramanian et al., 2021). Namely, of the 898 genes found to undergo changes in 3’ UTR isoform usage, 524 of those genes underwent 3’ UTR shortening (Subramanian et al., 2021). We thus focused on these genes that underwent 3’ UTR shortening.

In line with our observations of defective spermatogenesis, reduced sperm migration and defective oocyte morphology, gene ontology analysis of the 524 genes undergoing 3’ UTR shortening in *cfim-1(lf)* worms revealed enrichment of 57 genes with various germline and related developmental functions, including sex differentiation, ameboidal cell migration and oocyte development in *cfim-1(lf)* mutants (**Fig. 4A**, **Fig. S3**; **Table 1**). Comparing the difference in median proximal poly(A) site usage between *cfim-1* and wildtype worms (pPAU_MUT_ – pPAU_WT_), a measurement of 3’ UTR shortening, we found that this value was higher when examining patterns of only genes with fertility-related functions (0.260 versus 0.089 for all genes undergoing changes in 3’ UTR isoform usage; **Fig. 4A**). We compared bulk transcript abundance with 3’ UTR isoform usage and observed changes in 3’ UTR isoform usage of these sex-related genes occur without an appreciable change in overall transcript abundance between wildtype and *cfim-1(lf)* mutant worms (**Fig. 4B**, **Fig. S3**). Overall, these results suggest that the fertility defects in *cfim-1(lf)* worms are the result of changes in the APA landscape.

**Figure 4:**
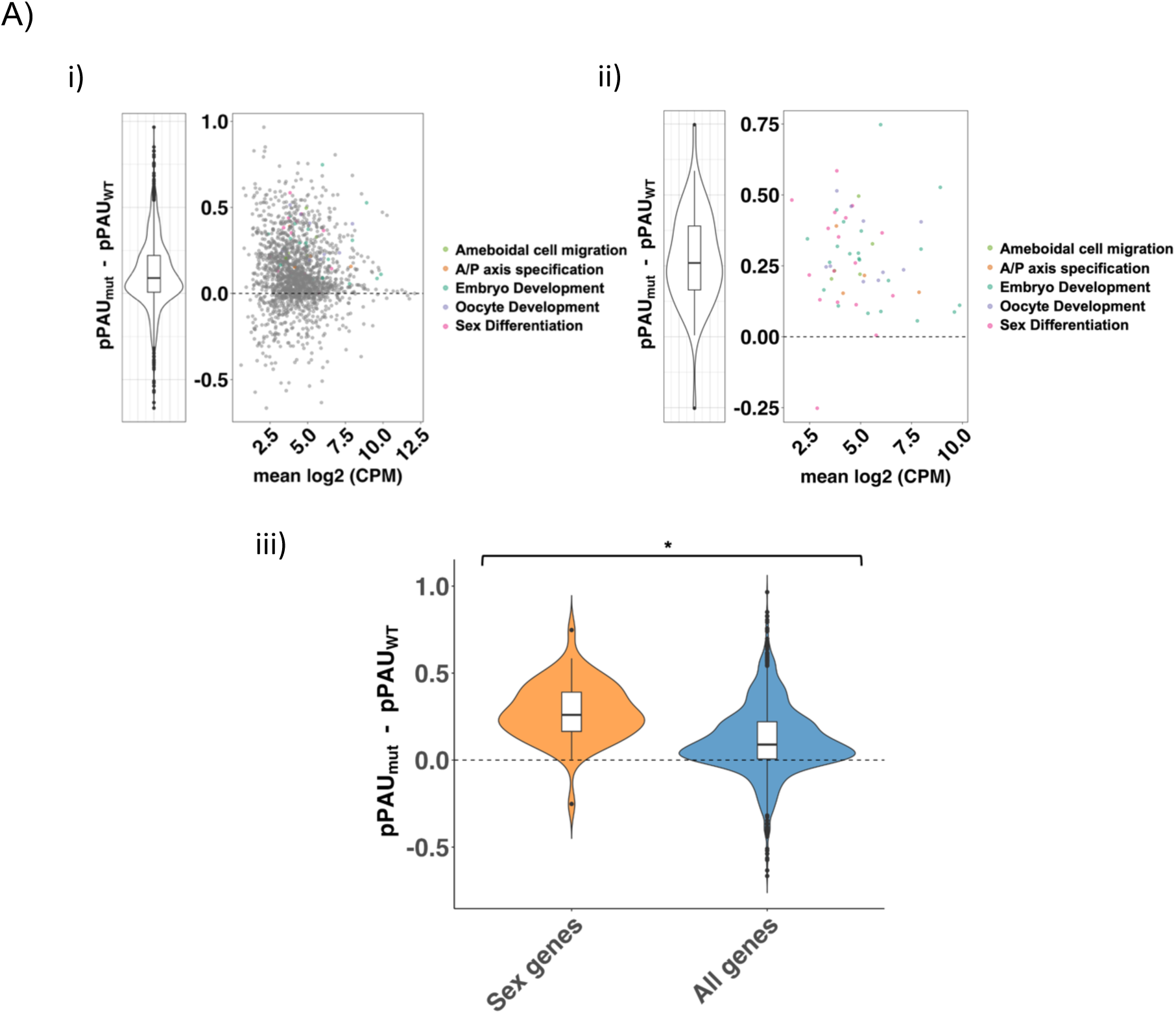
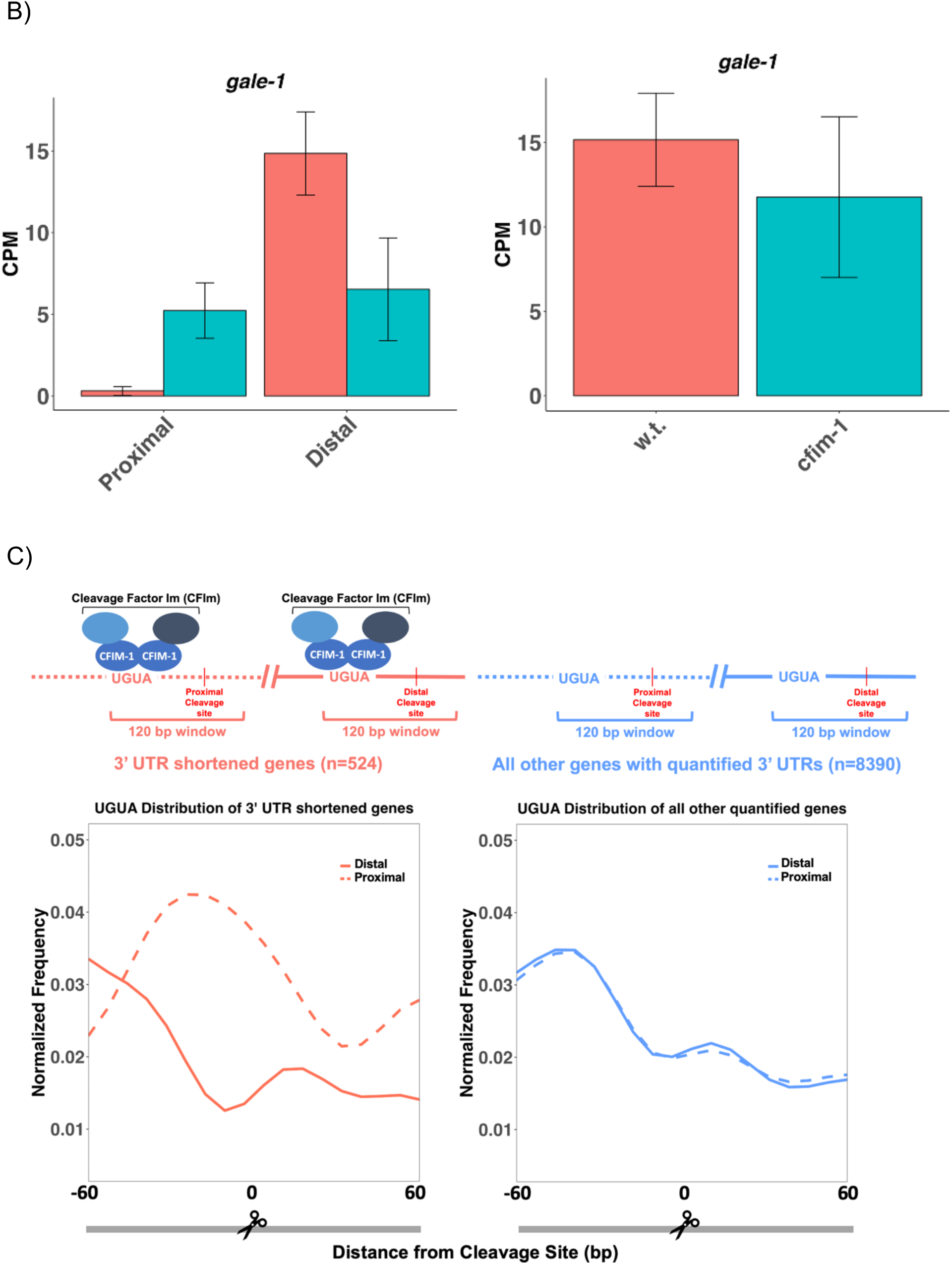
*cfim-1(lf)* elicits transcriptome-wide changes in alternative polyadenylation, including in genes with known functions in fertility. A) Comparisons of global patterns of APA in *cfim-1(lf)* worms. (i) Difference in proximal polyadenylation usage (pPAU) of all genes undergoing changes in APA in *cfim-1(lf)* (“pPAU_mut_”) versus wildtype worms (“pPAU_WT_”). Genes with known fertility and developmental roles (*n=57*) are highlighted. (ii) Same as (i) but for just for the highlighted gene set (*n=57*). (iii) Comparison of differences in proximal poly(A) site usage between *cfim-1* and wildtype worms (pPAU_MUT_ – pPAU_WT_) for genes with fertility-related functions and all genes undergoing APA. * p < 0.05 (Mann-Whitney test). **B)** Sample CPM plots showing isoform-specific (left) versus bulk changes (right) in RNA between wildtype and *cfim-1(lf)* mutants for the gene *gale-1*. Each bar represents the average of three replicates. For both graphs, red bars indicate expression in wildtype worms, while blue bars indicate expression in *cfim-1(lf)* mutants. **C)** Distribution of the canonical binding motif of the CFIm complex, UGUA, in the 120 bp window flanking the distal (solid lines) and proximal (dashed lines) cleavage sites of the genes undergoing 3’ UTR shortening in *cfim-1(lf)* worms (red) compared to all genes with quantified 3’ UTR isoforms.

**Table 1:** Description of the 57 fertility related genes subject to 3’ UTR shortening in *cfim-1(lf)* mutants.

| Symbol | Ensembl Gene ID | Chr | Description |
| --- | --- | --- | --- |
| kgb-1 | WBGENE00002187 | IV | GLH-binding kinase 1 |
| arx-4 | WBGENE00021170 | III | putative actin-related protein 2/3 complex subunit 2 |
| npp-8 | WBGENE00003794 | IV | Nuclear Pore complex Protein;Nucleoporin_C domain-containing protein;Nucleoporin_N domain-containing protein |
| ced-10 | WBGENE00000424 | IV | Ras-related protein ced-10 |
| smp-2 | WBGENE00004890 | I | Sema domain-containing protein |
| ceh-18 | WBGENE00000441 | X | Homeobox protein ceh-18;uncharacterized protein |
| mpk-1 | WBGENE00003401 | III | Mitogen-activated protein kinase mpk-1 |
| apc-11 | WBGENE00000145 | III | Anaphase-promoting complex subunit 11 |
| ubc-9 | WBGENE00006706 | IV | SUMO-conjugating enzyme UBC9 |
| let-502 | WBGENE00002694 | I | Rho-associated protein kinase let-502 |
| pal-1 | WBGENE00003912 | III | Homeobox protein pal-1 |
| pkc-3 | WBGENE00004034 | II | Protein kinase C-like 3 |
| arf-1 | WBGENE00000182 | III | ADP-ribosylation factor 1-like 2 |
| nop-1 | WBGENE00017774 | III | Pseudocleavage protein nop-1 |
| pak-1 | WBGENE00003911 | X | Serine/threonine-protein kinase pak-1 |
| mel-32 | WBGENE00003214 | III | Serine hydroxymethyltransferase |
| icd-1 | WBGENE00002045 | II | Transcription factor BTF3 homolog |
| gon-14 | WBGENE00002148 | V | Dimer_Tnp_hAT domain-containing protein |
| cye-1 | WBGENE00000871 | I | G1/S-specific cyclin-E |
| mek-1 | WBGENE00003185 | X | Dual specificity mitogen-activated protein kinase kinase mek-1 |
| wrt-1 | WBGENE00006947 | II | Warthog protein 1 N-product |
| capg-1 | WBGENE00009254 | I | Condensin complex subunit capg-1 |
| cki-2 | WBGENE00000517 | II | CKI family (Cyclin-dependent Kinase Inhibitor) |
| csr-1 | WBGENE00017641 | IV | Piwi domain-containing protein |
| sem-5 | WBGENE00004774 | X | Sex muscle abnormal protein 5 |
| hda-2 | WBGENE00001835 | II | Putative histone deacetylase 2 |
| wsp-1 | WBGENE00006957 | IV | WASP (actin cytoskeleton modulator) homolog |
| oma-1 | WBGENE00003864 | IV | CCCH-type zinc finger protein oma-1 |
| unc-129 | WBGENE00006852 | IV | TGF_BETA_2 domain-containing protein |
| mel-26 | WBGENE00003209 | I | Protein maternal effect lethal 26 |
| vha-17 | WBGENE00009882 | IV | V-type proton ATPase subunit e |
| skr-1 | WBGENE00004807 | I | Skp1-related protein |
| rbx-1 | WBGENE00004320 | V | RING-box protein 1 |
| cbp-1 | WBGENE00000366 | III | Protein cbp-1 |
| usp-46 | WBGENE00011216 | III | Ubiquitin carboxyl-terminal hydrolase 46 |
| gon-1 | WBGENE00001650 | IV | A disintegrin and metalloproteinase with thrombospondin motifs gon-1 |
| rad-51 | WBGENE00004297 | IV | DNA repair |
| arx-2 | WBGENE00000200 | V | Actin-related protein 2 |
| pro-2 | WBGENE00007413 | II | Nucleolar complex |
| cul-1 | WBGENE00000836 | III | Cullin-1 |
| sdn-1 | WBGENE00004749 | X | Syndecan;putative syndecan |
| cyb-1 | WBGENE00000865 | IV | G2/mitotic-specific cyclin-B1 |
| mix-1 | WBGENE00003367 | II | Mitotic chromosome and X-chromosome-associated protein mix-1 |
| pptr-2 | WBGENE00007554 | V | Serine/threonine protein phosphatase 2A regulatory subunit |
| math-33 | WBGENE00010406 | V | Ubiquitin carboxyl-terminal hydrolase 7 |
| mig-2 | WBGENE00003239 | X | Uncharacterized protein |
| par-5 | WBGENE00003920 | IV | 14-3-3-like protein 1;14_3_3 domain-containing protein |
| hmp-1 | WBGENE00001978 | V | Alpha-catenin-like protein hmp-1 |
| npp-3 | WBGENE00003789 | II | Nuclear Pore complex Protein |
| pph-4.1 | WBGENE00004085 | III | Serine/threonine-protein phosphatase 4 catalytic subunit 1 |
| gale-1 | WBGENE00008132 | I | UDP-glucose 4-epimerase |
| bub-3 | WBGENE00013209 | II | Mitotic checkpoint protein bub-3 |
| lev-11 | WBGENE00002978 | I | Tropomyosin isoforms a/b/d/f;Tropomyosin isoforms c/e;Uncharacterized protein |
| unc-112 | WBGENE00006836 | V | Protein unc-112 |
| dyn-1 | WBGENE00001130 | X | Dynamin |
| pptr-1 | WBGENE00012348 | V | Serine/threonine-protein phosphatase 2A<br>regulatory subunit pptr-1 |
| let-2 | WBGENE00002280 | X | Collagen alpha-2(IV) chain |

To gain more mechanistic insight, we wondered how CFIM-1 might influence cleavage site selection to dictate the APA landscape. In mammals, the CFIm complex, which includes the CFIM-1 homolog NUDT21, preferentially binds to its canonical binding motif, UGUA, upstream of the cleavage site, facilitating recruitment of the CPA machinery needed for 3’ end processing of pre-mRNAs (Brown and Gilmartin, 2003; Venkataraman et al., 2005; Yang et al., 2010; Yang et al., 2011; Martin et al., 2012; Brumbaugh et al., 2018). This binding occurs selectively at distal cleavage sites of these genes, so that its depletion causes preferential utilization of proximal cleavage sites.

This is corroborated by the finding that this UGUA motif is enriched upstream of distal cleavage sites of 3’ UTR shortened genes in *NUDT21* depleted mammalian cells (Brumbaugh et al., 2018). We sought to determine if the same was true in worms.

Noting the high sequence and predicted structural homology of CFIM-1 to NUDT21 (**Fig. S4**), we examined the distribution of the UGUA nucleotide sequence in the region flanking the cleavage sites of genes undergoing 3’ UTR shortening in response to abrogation of *cfim-1*. To our surprise, we found an enrichment of this motif at proximal, rather than distal, cleavage sites (**Fig. 4C, D**). This suggests a different mode of regulation in the worm, whereby recruitment of the CFIm complex may inhibit cleavage at these proximal sites (**Fig. 5**).

**Figure 5:**
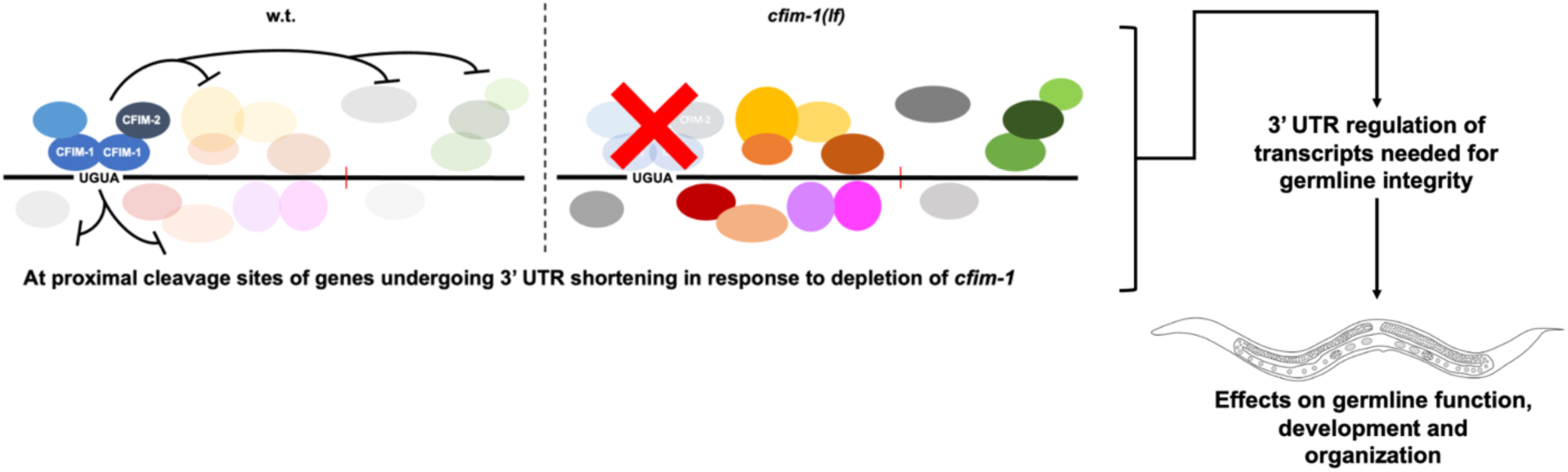
Cartoon model of how *cfim-1* might mediate its effects on post-transcriptional gene regulation and the associated effects on germline integrity.

## DISCUSSION

### *cfim-1* is crucial in post-embryonic germline integrity

It is well known that the core ensemble of cleavage and polyadenylation factors determine differential 3’ UTR isoform usage due to their effects on pre-mRNA processing. However, their post-embryonic functions, if any, have remained nebulous due their essential roles in embryonic development (Koscielny et al., 2014). Among these factors, mammalian *NUDT21* has been a recurrent focus of investigation due to its associated role in a variety of diseases, including cancer (Masamha et al., 2014; Sun et al., 2017; Tan et al., 2018; Chu et al., 2019; Yang et al., 2020; Xing et al., 2021; Subramanian et al., 2021). We previously reported a post-embryonic role in *C. elegans* vulva development for the worm *NUDT21* homolog, *cfim-1* (Subramanian et al., 2021). However, this effect was only observed in the context of hyperactivated LET-60/Ras signalling. Here, we report a fully-penetrant, temperature-dependent role for *cfim-1* in germline development and function. Intriguingly, after surveying an array of CPA factors that represent the core CPA machinery, we found that this temperature-dependent sterile phenotype was unique to *cfim-1* (**Fig. S2A-D**). For example, ablation of its canonical binding partner *cfim-2* resulted in sterility or markedly reduced brood size at all temperatures assayed (**Fig. S2C**). Thus, *cfim-1* is dispensable at the normal cultivation temperatures of 16-20 °C, but at 25 °C germline defects that result in sterility are observed. Noting the functioning of its ortholog NUDT21 as part of the multimeric CFIm protein complex in mammals, as well as its sequence and predicted structural homology to NUDT21 (**Fig. S4**), the uniqueness in terms of the dispensability of *cfim-1* in *C. elegans* may reflect differences in its stoichiometric requirements for facilitating cleavage and polyadenylation (Yang et al., 2010; Yang et al., 2010; Brumbaugh et al., 2018).

### Complex interplay among genes associated with development of the germline

Characterization of *cfim-1* mutants revealed multiple germline phenotypes. As cell cycle progression, mRNA localization and translation are spatially demarcated in the germline (West et al., 2018; Diag et al., 2018), we first quantified the distribution and abundance of germ cell nuclei and found that a reproducible hallmark of *cfim-1(lf)* mutants was ectopic accumulation of germ cell nuclei in the distal region (**Fig. 1B, E**). Given the developmental timing for the onset of sterility, which is concordant with the switch between oogenesis and spermatogenesis, we evaluated the quantity and quality of sperm produced. *cfim-1(lf)* mutants produce fewer sperm that are also aberrantly arranged along the gonadal arm, suggesting defects in gamete specification and motility (**Fig. 2C, D**). Furthermore, we found that *cfim-1(lf); spe-11::mScarlet* males fail to inseminate wildtype or *cfim-1(lf)* hermaphrodites at 25 °C. This occurs despite repeated copulation attempts by the *cfim-1(lf)* male worms. While we cannot discount the idea that failed insemination is due to blockage of the male reproductive tract, our overall findings suggest gross morphological defects and reduced motility of *cfim-1(lf)* sperm (**Fig. 2C, D**).

We also determined whether these sperm defects could solely explain the sterile phenotype by mating sterile *cfim-1(lf)* mutants to wildtype males at 25 °C. The failure to obtain viable outcross progeny, and the presence of morphologically abnormal oocytes (**Fig. S2B**), suggests multiple roles for *cfim-1* in the germline that likely culminate in sterility. The behaviour of wildtype sperm in the proximal reproductive tract of *cfim-1(lf)* worms reared at 25 °C revealed that even in the presence of wildtype sperm, fertilization fails to occur. This suggests that the morphological defects in oocytes may prevent them from being fertilized (**Video 1; Fig. 3B**).

The complex germline phenotypes of *cfim-1(lf)* mutants are not surprising, given that it is involved in processing and regulating hundreds genes, many of which are expressed in the germline, via tandem alternative polyadenylation (Diag et al., 2018; West et al., 2018; Subramanian et al., 2021). We attempted to identify suppressors of the temperature-dependent sterile phenotype by screening 18,900 mutagenized genomes. Our inability to identify genetic suppressors of *cfim-1* mutants suggest that the *cfim-1* germline phenotype is due to its effects on multiple genes that regulate germline integrity and function. Teasing this apart will require systematic functional analysis of individual genes with germline functions undergoing alterations in their APA patterns in response to loss of *cfim-1* (**Table 1**).

### An inhibitory model for *cfim-1* mediated alternative polyadenylation

Previous work on the mechanism by which NUDT21, the mammalian homolog of CFIM- 1, mediated 3’ UTR site selection suggested that inhibition of the CFIm complex biases pre-mRNA processing by the core CPA machinery towards usage of proximal 3’ UTR cleavage sites (Brumbaugh et al., 2018). Namely, the CFIm complex binds preferentially upstream of distal cleavage sites in the 3’ UTRs of pre-mRNAs to facilitate their cleavage and polyadenylation. In the absence of the CFIm complex, the core CPA machinery binds to proximal cleavage sites to mediate post-transcriptional processing.

This model was supported, in part, by the finding that the UGUA tetrameric sequence, the canonical binding motif of the CFIm complex, is preferentially enriched just upstream of the distal cleavage site of genes subject to APA changes following *NUDT21* knockdown in mouse embryonic fibroblasts (Brumbaugh et al., 2018). Noting that CFIM-1 has high sequence and predicted structural homology to mammalian NUDT21, we also examined the enrichment of this motif in the worm (Jumper et al., 2021; Yang et al., 2011; **Fig. 4C**, **S4**). Genes subject to increased usage of proximal 3’ UTRs in the absence of *cfim-1* were enriched for the UGUA motif upstream of the proximal 3’ UTRs, the opposite of what is observed in mammals. This points to the idea that CFIM-1 may actually inhibit usage of proximal cleavage sites at 3’ UTRs in *C. elegans*. The suggestion of pleiotropic effects of *cfim-1* mediated regulation of APA across metazoans is intriguing, but more biochemical work will be needed to elucidate how CFIM-1 and its homologs regulate alternative polyadenylation.

## MATERIALS AND METHODS

### Brood Counts

*Caenorhabditis elegans* worms of the desired strains were synchronized at the L1 stage of development by the bleaching protocol. Briefly, gravid non-starved worms were washed off from crowded, but non-starved, 6 cm NGM plates and resuspended in 1 ml of one day old hypochlorite solution (550 μl MilliQ Water, 250 μl 1M NaOH, 200 μl commercial bleach). Worms were manually agitated until cuticles had just begun to dissolve. Worm carcasses and eggs were immediately pelleted by spinning at 4000 RPM for 30 seconds. The supernatant was removed, and worms were resuspended in 1 ml of fresh M9 solution, followed by subsequent centrifugation at 4000 RPM for 30 seconds. This process of washing the worms in M9 was repeated five times. After the final resuspension in M9 buffer, eggs were allowed to hatch with gentle agitation overnight (12 hours), producing a life-stage synchronized population of L1 worms. These synchronized worms were reared from the larval L1 stage (1 worm/plate) at the desired conditions and temperature. Once they reached the adult stage, worms were transferred daily until they no longer laid eggs, and their F1 progeny counted using a dissecting microscope.

### Temperature Shift Experiments

Life-staged synchronized worms reared from L1, obtained as described above, were shifted at timepoints specified in figures and assessed for brood size.

### Germline image analysis

Germlines of transgenic strains expressing *rec- 8::mNeonGreen* and *him-3::mScarlet* were volumetrically imaged on a Leica SP8 Lightsheet Confocal microscope. Germ cell nuclei were manually annotated from the distal end to the gonadal bend to obtain quantifications and 3D positional information (Venkatachalum et al., 2016). We defined all mitotic germ cell nuclei as those that are positive for mNeonGreen but not for mScarlet, given the ubiquity of *rec-8::mNeonGreen* expression in germ cell nuclei. We defined all meiotic nuclei as those positive for the mScarlet signal. For germline linearization, we used a modified version of a published framework for characterizing germ nuclei distribution along the gonad arm (Toraason et al., 2021). In brief, annotated germlines are manually segmented in a linear fashion along their DP axis and projected onto a 2D plane. Next, each germ nucleus is linked to its nearest neighbour line segment, which were then aggregated and normalized onto a single one-dimensional axis. This facilitates a cross comparison of their densities based on their position along the DP gonad arm.

### CRISPR/Cas9 Gene Editing

All transgenic *C. elegans* used in this study were generated using a modified CRISPR/Cas9 based protocol (Eroglu et al., 2023). Cas9- crRNA-tracrRNA RNP complexes, along with the corresponding single-stranded DNA repair templates, were administered via injection into mature wildtype N2 hermaphrodite worms to facilitate the incorporation of the intended genetic lesion. A co-injection marker for *dpy-10* was used to identify candidate crispants. Successful crispants were identified by PCR and sequencing and outcrossed once to N2 males to remove the co-injection marker.

### Sperm Motility Assays

We employed a modified version of a sperm motility assay to quantify and assess sperm trajectories (Hu et al., 2019). Transgenic males of the desired genotype that expressed *spe-11::mScarlet* were mated to either wildtype (N2) or *cfim-1(lf)* hermaphrodites at 25 °C. Hermaphrodite worms were immobilized in 1%/0.1% w/v Tricaine/Tetramisole solution, mounted on 4% agarose pads and immediately imaged by timelapse microscopy on a Leica DMRA compound microscope. Each germline was imaged for 15 minutes at a capture rate of one fluorescent image every 15 seconds. Before and immediately following acquisition of fluorescent images, DIC images were acquired to determine relative worm position to confirm that worm body position had not changed during timelapse imaging. Sperm were manually tracked using TrackMate to define path lengths, measured in pixels travelled per unit time. These values were used to calculate vector velocity relative to the spermatheca and path-length speed (Ershov et al., 2022).

### RNA-sequencing analysis

We previously conducted RNA sequencing on transgenic strains: on9[GFP::3xFlag::*cfim-1*-8bp dele] I strain and reference wildtype (N2) worms, which allowed us to measure changes in the abundance of 3′ UTR isoforms (Subramanian et al., 2021). From the RNA sequencing data, genes were categorized based on changes in 3’ UTR transcript isoform levels between N2 and *cfim-1(lf)* worms (Subramanian et al., 2021). One of these groups consisted of genes in *cfim-1(lf)* worms that showed 3’ UTR shortening (which we denote as “3’ UTR shortened genes” in the figures). This set of genes (*n=524*) demonstrated a shift towards higher abundance of the shortened 3′ UTR transcript isoform compared to the elongated 3′ UTR isoform in the *cfim-1(lf)* worms. From this set of genes, we performed gene ontology analysis to define enriched pathways and biological functions (Thomas et al., 2022). Using as our background all genes that were quantified in our analyses, we identified enrichment in 57 genes with sex and related developmental functions, including ameboidal cell migration (*n=5*), anterior-posterior axis specification (*n=7*), embryo development (*n=27*), oocyte development (*n=10*) and sex differentiation (*n=18*).

### Sequence enrichment analysis

From the 524 genes we classified as undergoing 3’ UTR shortening, we also calculated the incidence of the UGUA motif, the canonical binding sequence of the CFIm complex, in the 120 bp region flanking the poly(A) site for each of these genes. We chose to analyze this region as the median distance between the to the poly(A) site element, where most core cleavage and polyadenylation factors bind, and the cleavage site was previously shown to be 19 bp (Steber et. al, 2019). We extended this region by a 50 bp window both upstream and downstream to account for CFIM-1, as part of the CFIm complex, which may engage these sites slightly further away. We created 2 matrices, representing the regions surrounding the distal and proximal cleavage sites, to numerically represent the incidence and position of these motifs. For aggregate analysis, each matrix was condensed by summing columns. The sums were then normalized to the number of genes. The resulting frequencies were grouped into 20 bins, with subsequent normalization for and smooth curve fitting for graphing, in accordance with previously described methods (Brumbaugh et al., 2018). **Creation of figures:** All graphs and videos were generated in RStudio, Biorender, ChimeraX or Fiji.

## ACKNOWLEDGEMENTS

We are grateful to John Calarco for his insightful comments on this manuscript. Confocal microscopy was conducted using the Leica SP8 Imaging Facility at The Hospital for Sick Children in Toronto, Canada. Some *C. elegans* strains were obtained from the Caenorhabditis Genetics Center (CGC), which receives support from the NIH Office of Research Infrastructure Programs (P40 OD010440). This research was funded by the Canadian Institutes of Health Research (CIHR, grant PJT 155928). W.B.D. holds the Canada Research Chair in Animal Models of Human Disease.

---

**Figure S1:**
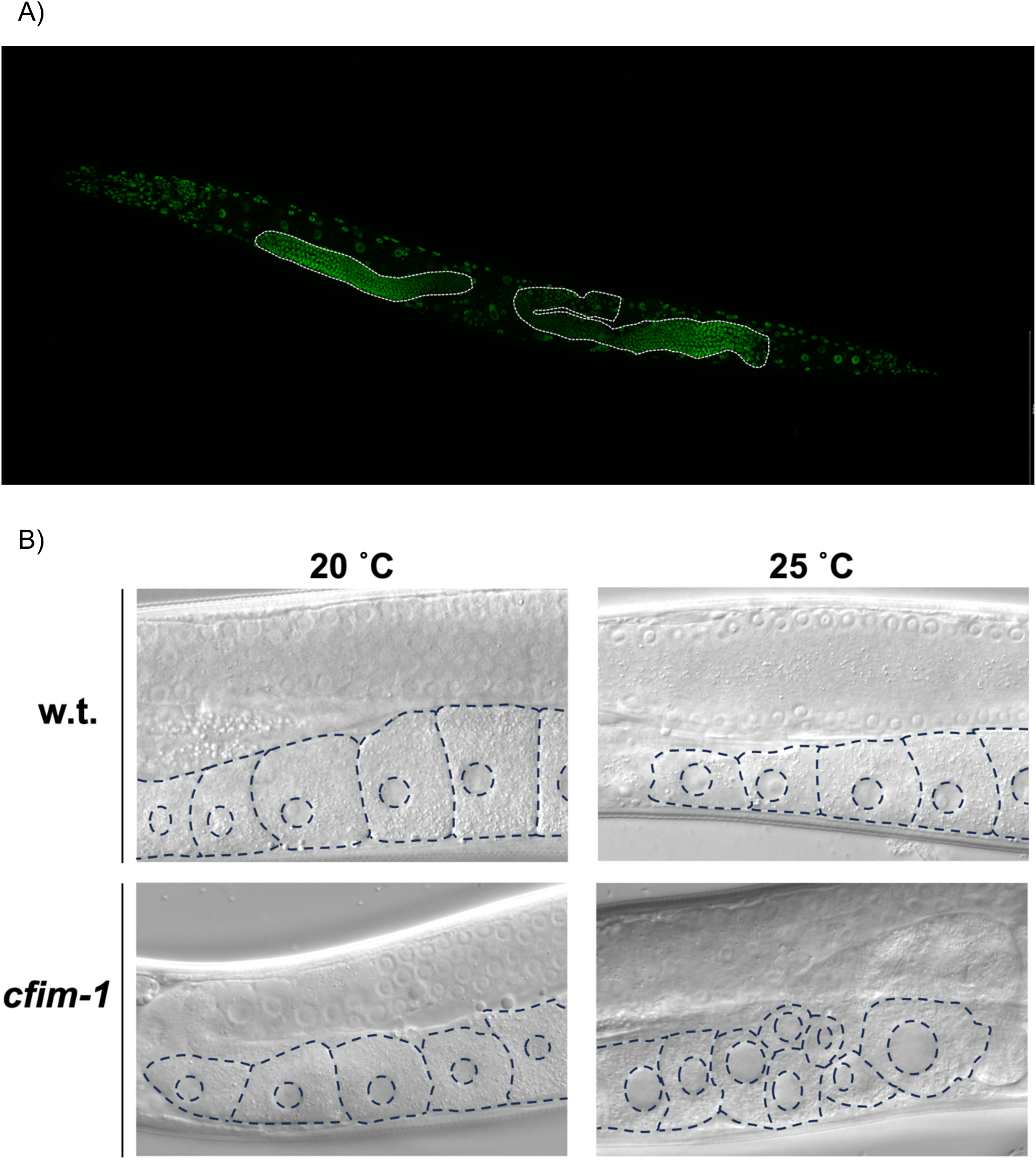
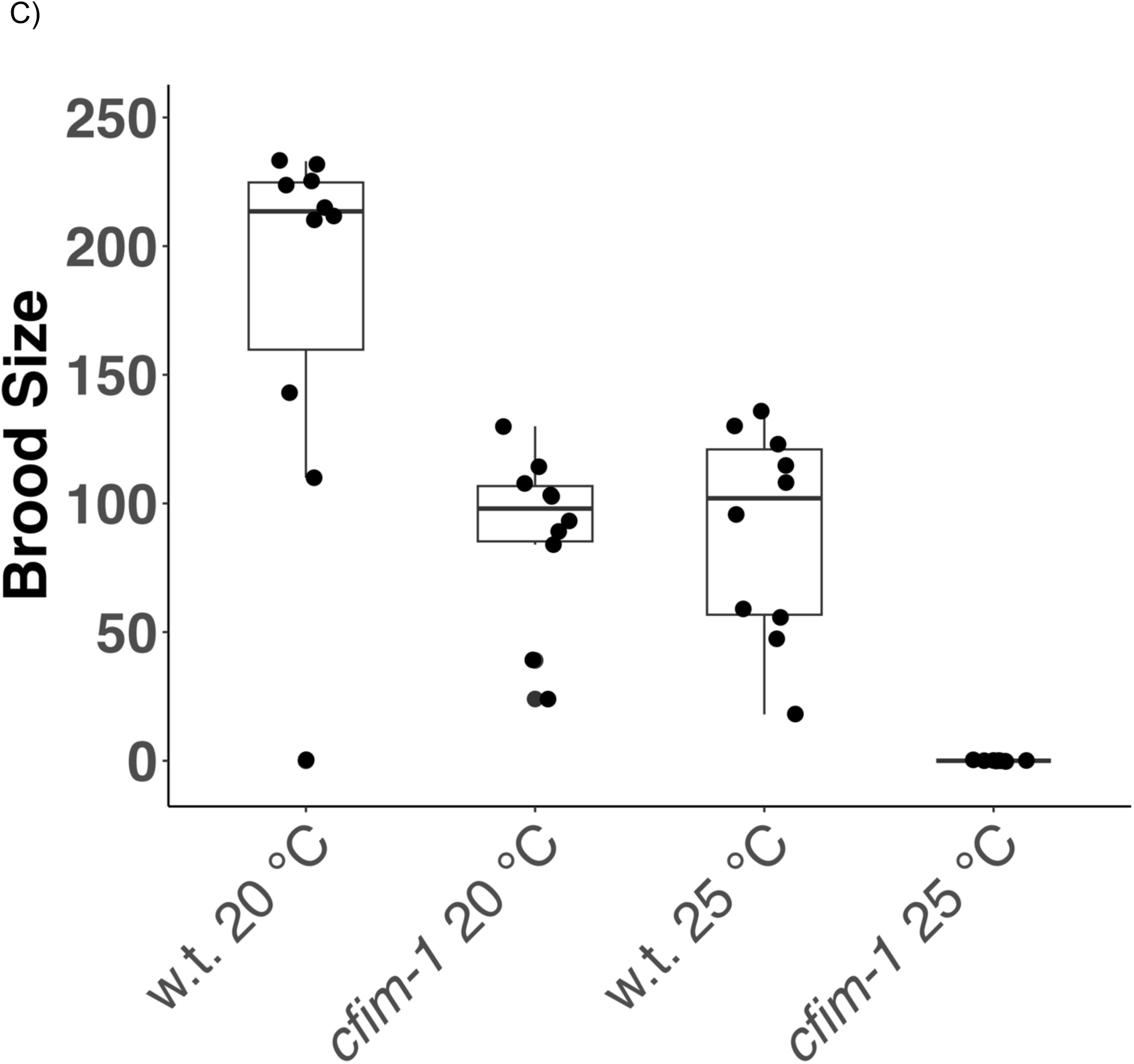
Analysis of germline phenotypes of *cfim-1(lf)* worms. **A)** Representative pseudo colored maximum intensity projection fluorescence image depicting whole body expression of CFIM-1. Germline arms are outlined. **B)** Proximal germline arm of *cfim- 1(lf)* worms showing oocyte distribution and morphology. **C)** Brood counts for a full in-frame deletion of the *cfim-1* locus and N2 wildtype (“w.t.”) worms at the indicated temperatures. Each dot represents the brood size for one worm. Box and whisker overlays show the median with the first and third quartiles. n=11 for wildtype 20 °C, n=12 for *cfim-1* 20 °C, n=10 for wildtype 25 °C and n=10 for *cfim-1 25 °C*.

**Figure S2:**
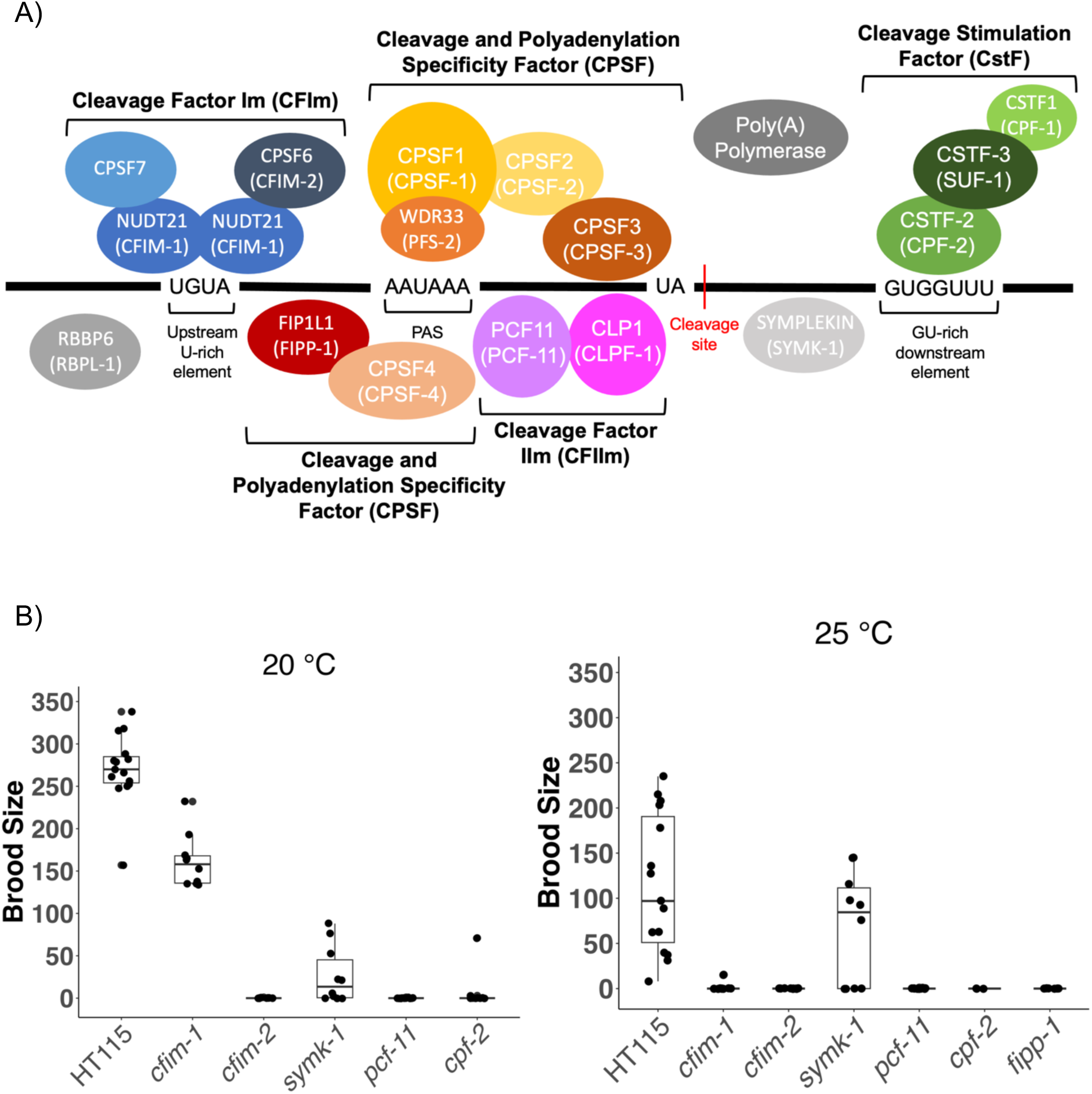
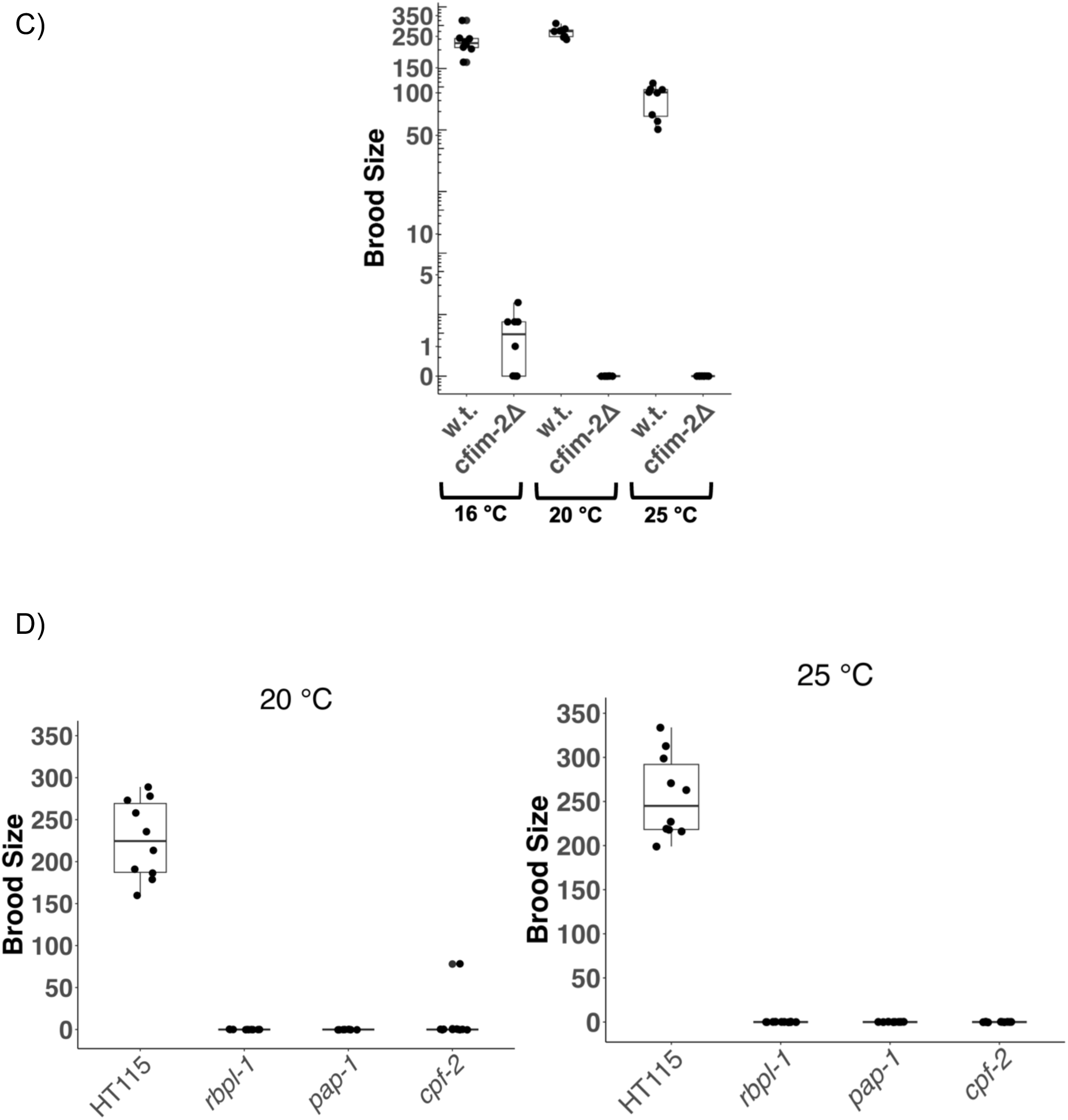
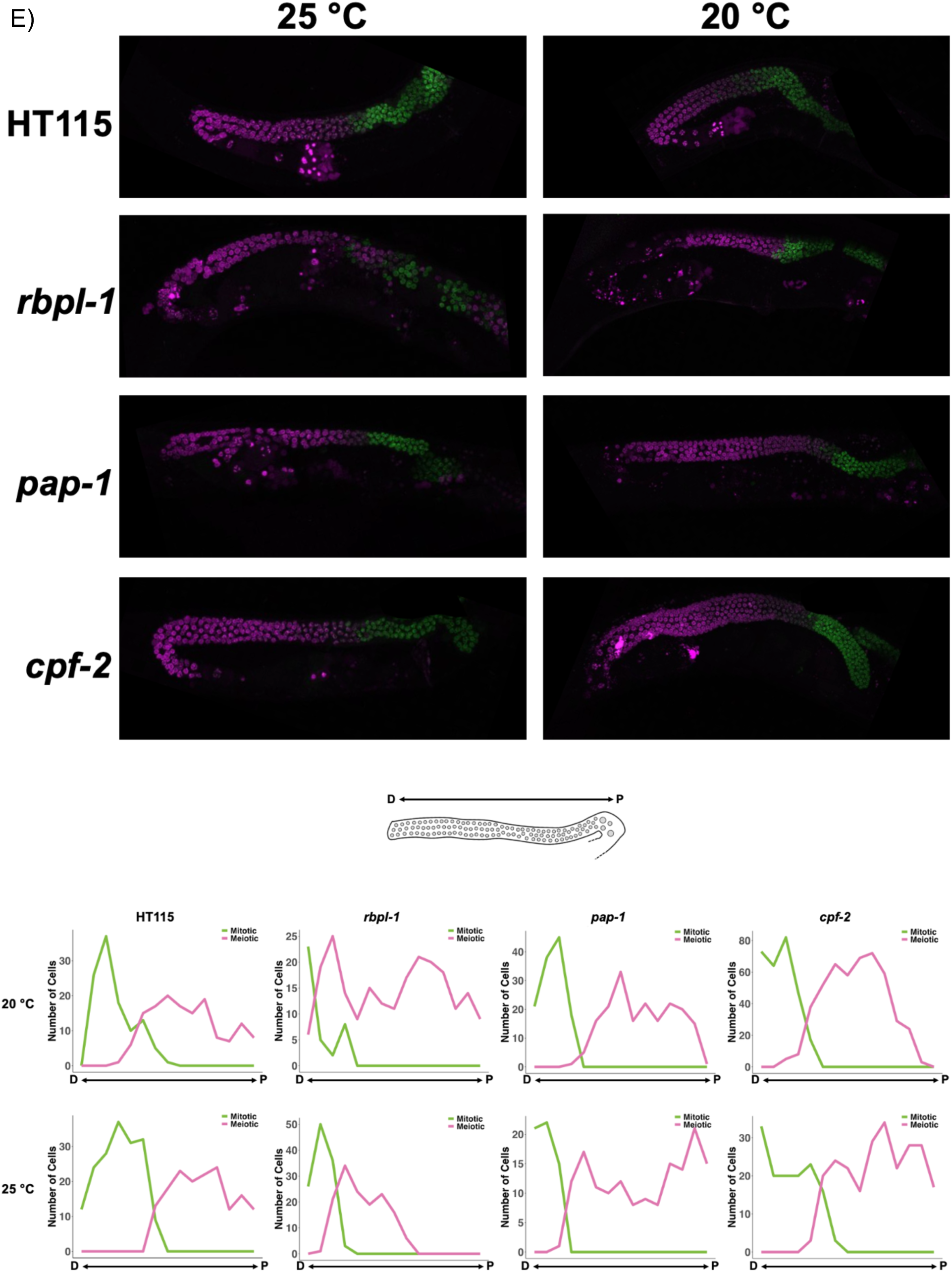
Characterization of fecundity and germline phenotypes across core CPA factors. **A)** Cartoon model of proposed cleavage and polyadenylation complex and associated machinery, adapted from Steber et al., 2019. **B)** Brood counts for several core cleavage and polyadenylation factors subject to RNAi at the indicated temperatures. **C)** Brood counts for a full in-frame deletion of the *cfim-2* locus. **D)** Brood counts for worms subjected to RNAi against indicated core CPA factors. **E)** Germline analysis for core CPA factors assayed in Fig. S2D. Top panels depict maximum intensity projections of *rec-8::mNeonGreen* (green) and *him-3::mScarlet* (magenta) for indicated RNAi conditions. Bottom panels show representative germ cell nuclei distribution plots, with green lines indicating mitotic germ cell nuclei and magenta lines indicating meiotic germ cell nuclei.

**Figure S3:**
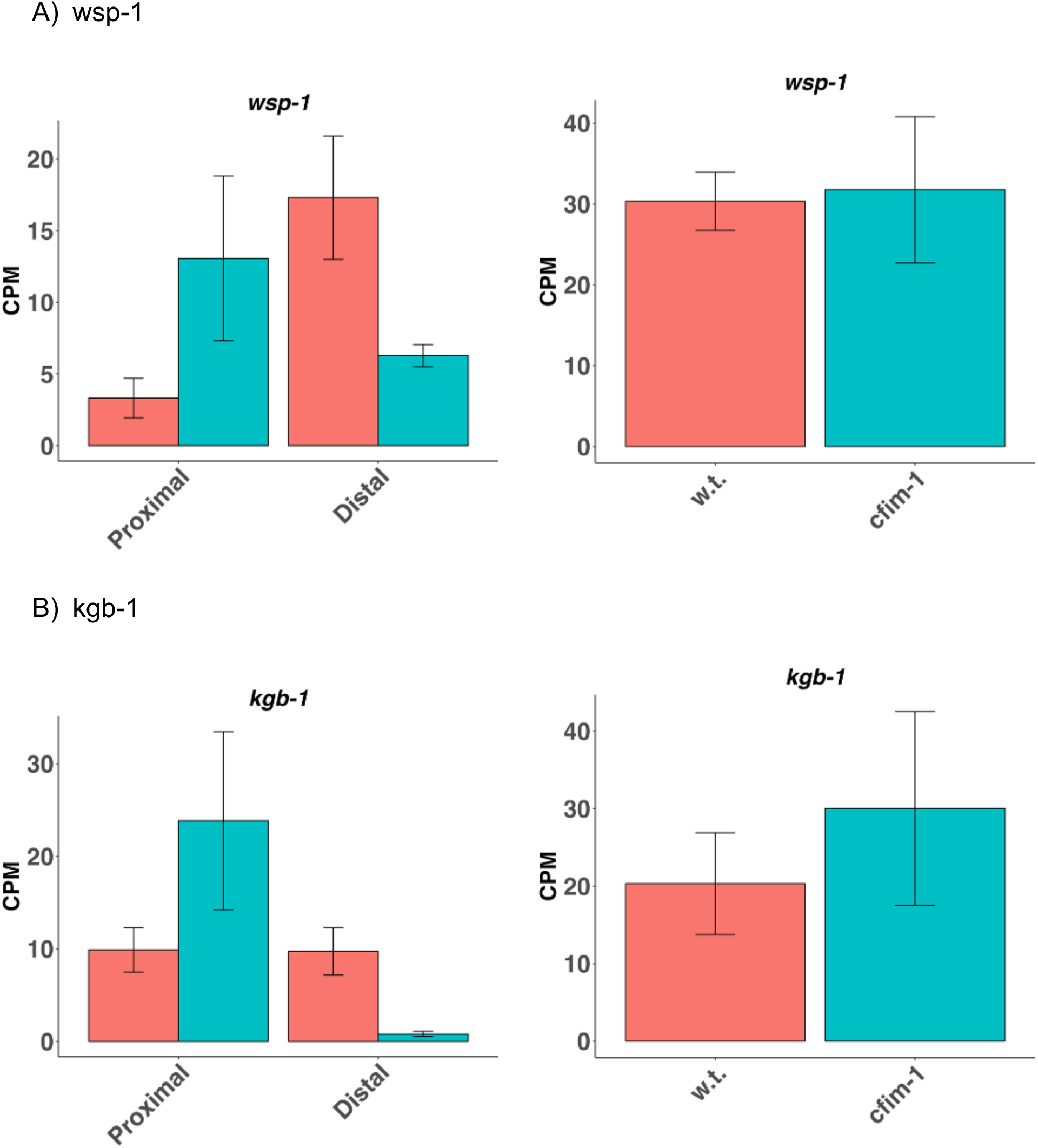
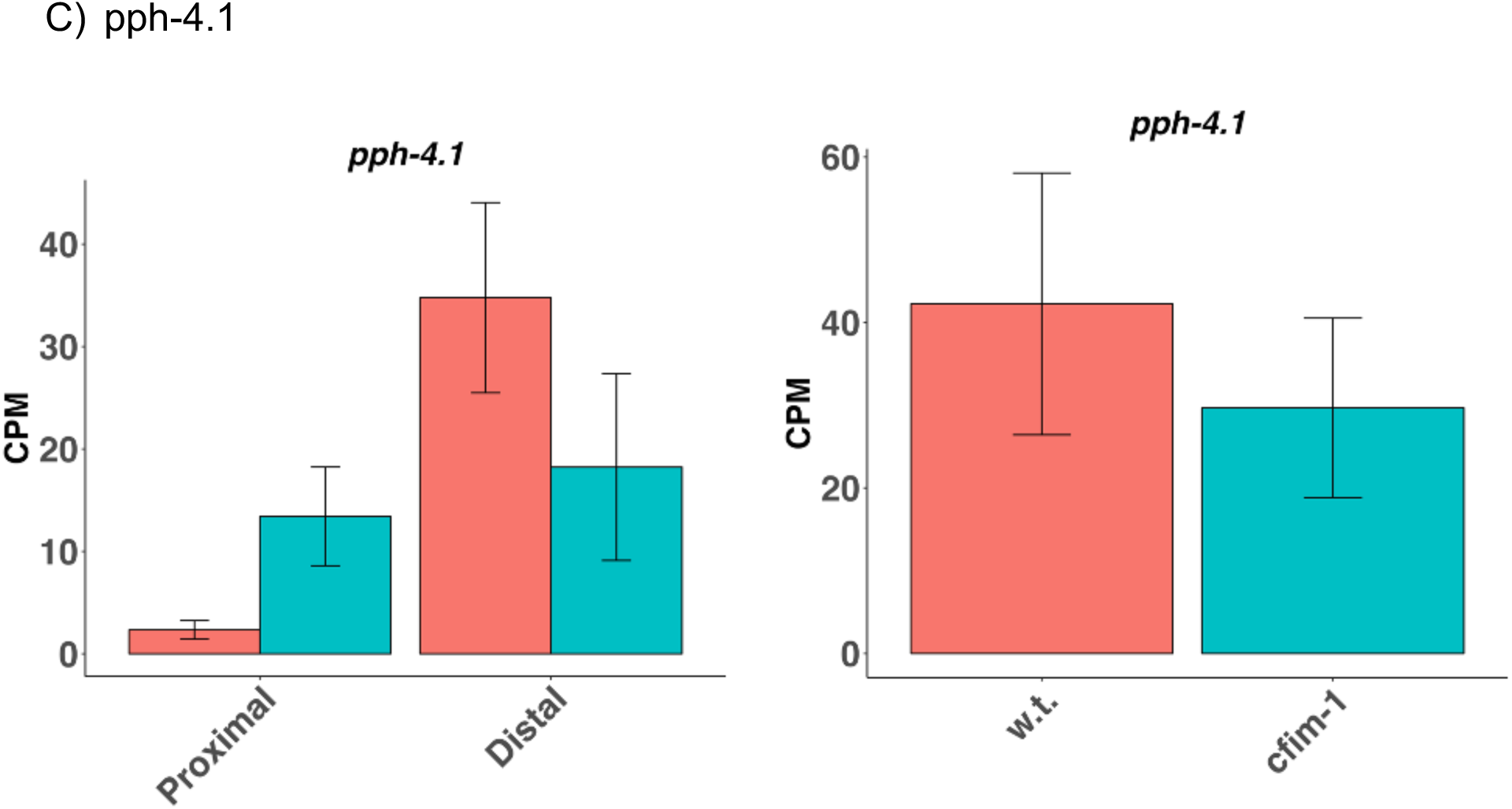
Loss-of-function of *cfim-1* induces changes in alternative polyadenylation, including in genes with known male and female germline functions. Representative changes in 3’ UTR isoform abundance (graphs at left), without appreciable changes in bulk RNA (graphs at right) for sample genes with functions relating to **A)** ameboidal cell migration, **B)** oocyte development and **C)** embryo development, upon abrogation of *cfim-1*. Bars indicate median CPM of sequenced replicates for each condition. For graphs at left, red bars indicate changes in wildtype worms, while blue bars indicate changes in *cfim-1(lf)* mutants. Error bars indicate standard deviation for the isoform or transcript of the indicated condition.

**Figure S4:**
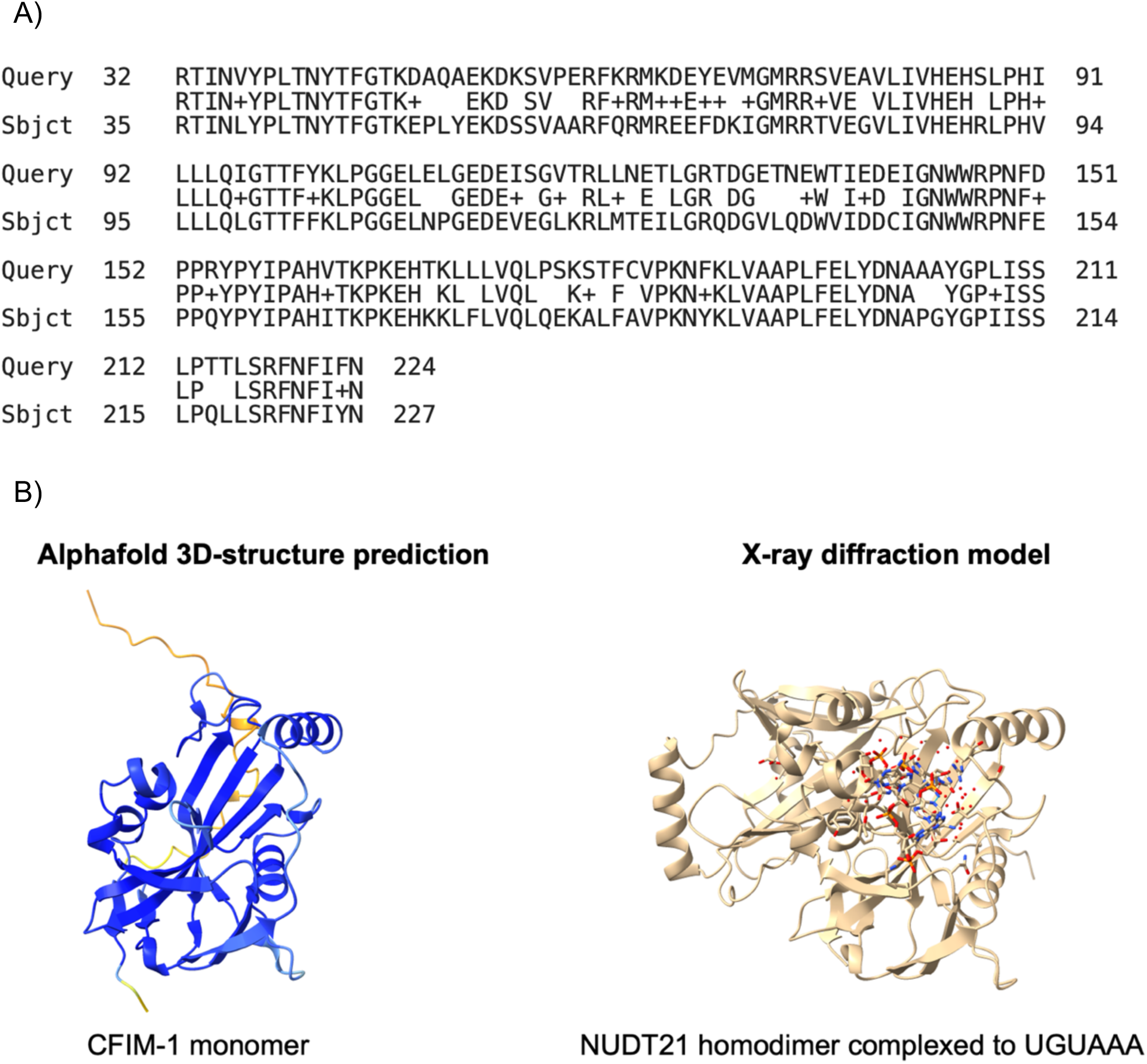
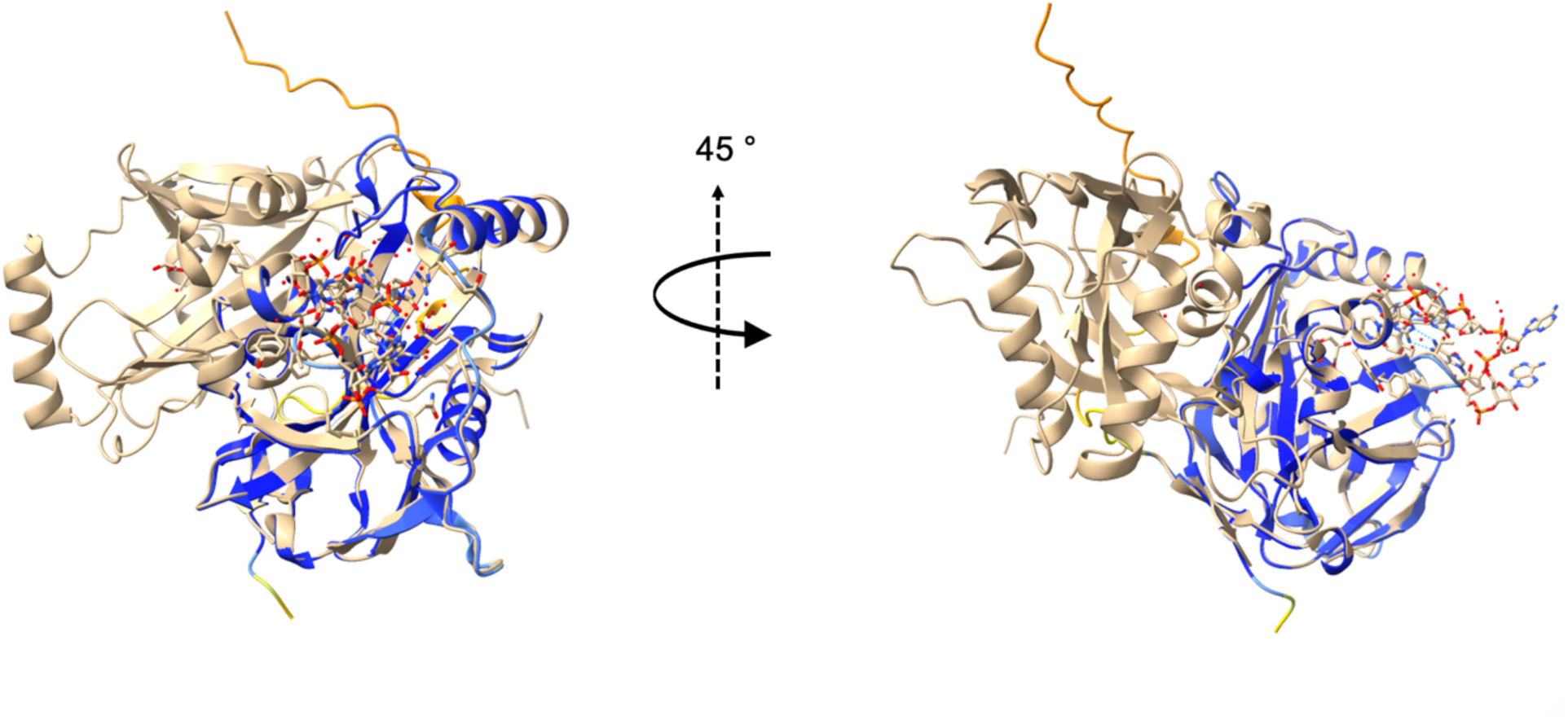
Comparison of sequence and predicted structural homology between NUDT21 and CFIM-1. **A)** Pairwise amino acid sequence alignment of *cfim-1* (Query) and NUDT21 (Subject). **B)** Structural homology between the *NUDT21* homodimer (beige) bound to UGUAAA, obtained by X-ray crystallography, and a predicted model of the *cfim-1* monomer (blue)

**Video 1:** Timelapse movie of sperm motility of *spe-11::mScarlet* males mated to *cfim- 1(lf)* worms at 25 °C. Movie represents 15 minutes of sperm motility, and has been sped up 217x. Beginning and end frames define relative gonad position.

**Video 2:** Timelapse movie of sperm motility of *spe-11::mScarlet* males mated to N2 worms at 25 °C. Movie represents 15 minutes of sperm motility, and has been sped up 217x. Beginning and end frames define relative gonad position.

Videos are available at: https://figshare.com/s/41ee75c661cb38b86bb4

